# The ER membrane protein complex governs lysosomal turnover of a mitochondrial tail-anchored protein, BNIP3, to restrict mitophagy

**DOI:** 10.1101/2023.03.22.533681

**Authors:** Jose M Delgado, Logan Wallace Shepard, Sarah W Lamson, Samantha L Liu, Christopher J Shoemaker

**Author notes:** (to C.J.S).

## Abstract

Lysosomal degradation of autophagy receptors is a common proxy for selective autophagy. However, we find that two established mitophagy receptors, BNIP3 and BNIP3L/NIX, violate this assumption. Rather, BNIP3 and NIX are constitutively delivered to lysosomes in an autophagy-independent manner. This alternative lysosomal delivery of BNIP3 accounts for nearly all of its lysosome-mediated degradation, even upon mitophagy induction. To identify how BNIP3, a tail-anchored protein in the outer mitochondrial membrane, is delivered to lysosomes, we performed a genome-wide CRISPR screen for factors influencing BNIP3 flux. By this approach, we revealed both known modifiers of BNIP3 stability as well as a pronounced reliance on endolysosomal components, including the ER membrane protein complex (EMC). Importantly, the endolysosomal system regulates BNIP3 alongside, but independent of, the ubiquitin-proteosome system (UPS). Perturbation of either mechanism is sufficient to modulate BNIP3-associated mitophagy and affect underlying cellular physiology. In short, while BNIP3 can be cleared by parallel and partially compensatory quality control pathways, non-autophagic lysosomal degradation of BNIP3 is a strong post-translational modifier of BNIP3 function. More broadly, these data reveal an unanticipated connection between mitophagy and TA protein quality control, wherein the endolysosomal system provides a critical axis for regulating cellular metabolism. Moreover, these findings extend recent models for tail-anchored protein quality control and install endosomal trafficking and lysosomal degradation in the canon of pathways that ensure tight regulation of endogenous TA protein localization.

## INTRODUCTION

Autophagy is an intracellular degradative pathway that clears unwanted cytoplasmic components such as damaged or superfluous organelles^1^. During autophagy, a unique double-membrane vesicle–the autophagosome–is generated around cargo. The completed autophagosome subsequently traffics to the lysosome where its content is degraded. The recognition and clearance of mitochondria by autophagy (hereafter mitophagy) are broadly implicated in aging, development, and disease^2, 3^. Immense progress has been made toward understanding the canonical PINK1/Parkin-dependent mitophagy pathway^2^. However, mitophagy can occur independently of this machinery (i.e., PINK1/Parkin-independent) where it is executed by less-understood mechanisms varying across cell type and physiological context^3^.

BNIP3 and BNIP3L/NIX are paralogous membrane proteins found on the outer mitochondrial membrane (OMM)^4, 5^. As mitophagy receptors, BNIP3 and NIX recruit key autophagy proteins, in particular the Atg8-family of proteins (LC3 and GABARAP families in humans), to the surface of targeted mitochondria^6–8^. Such interactions enforce cargo specificity by keeping the expanding autophagosomal membrane in close apposition to the targeted mitochondrion. The potency of these interactions is reflected in the observation that ectopic expression of BNIP3 or NIX is sufficient to induce selective mitophagy^9–11^. Thus, the expression and/or activation of BNIP3 and NIX must be appropriately constrained *in vivo* to spatiotemporally restrict aberrant mitophagy induction. Early studies identified transcriptional regulation by hypoxia-inducible factor 1 (HIF-1) as a key facet of BNIP3 and NIX regulation^4^. Consistent with this model, both BNIP3 and NIX expression and associated mitophagy are potently induced upon hypoxia onset^14^. Recently, multiple groups have extended this model, reporting that the ubiquitin-proteasome system (UPS) potently restricts BNIP3 and NIX levels to further curb mitophagy^12–17^. In light of these concepts, it is important to develop a unified understanding of how steady-state levels of these mitophagy receptors are established and maintained, and how this regulation governs underlying cell physiology.

BNIP3 and NIX are targeted to the OMM by a single, C-terminal transmembrane domain (TMD)^18^. This topology defines a diverse class of membrane proteins (∼50 in yeast, >300 in humans) known as tail-anchor (TA) proteins, which rely exclusively on post-translational insertion mechanisms^19–21^. TA protein targeting poses a fundamental and innate challenge for cells. The hydrophobicity of a TA TMD is a primary determinant of its localization, with mitochondrially-targeted TMDs having a lower hydrophobicity, on average, than those targeted to the ER^19, 22^. However, this relationship is not absolute. In the OMM, TA proteins are inserted via MTCH1/MTCH2, while mislocalized or aberrant TA proteins are extracted by ATAD1 (Msp1 in yeast)^23, 24^. In the ER membrane, TA proteins are inserted by either the ‘guided entry of TA proteins’ (GET) pathway or the ‘ER membrane protein complex’ (EMC), while mislocalized or aberrant TA proteins are extracted by ATP13A1 (Spf1 in yeast)^25–27^. Far from futile, dynamic cycles of TA protein insertion and extraction play a critical role in properly partitioning TA proteins despite limited and overlapping targeting information^28–33^. As representative TA proteins, BNIP3 and NIX are primarily localized to the OMM but have been demonstrated to localize to other membranes^34^. Consequently, exploration of BNIP3 and NIX regulation has the potential to reveal additional insights into TA protein quality control mechanisms.

Here we utilized a triple-negative breast cancer cell line MDA-MB-231, that forms dense hypoxic tumors *in vivo*, to study the post-translational regulation of BNIP3 in hypoxic and non-hypoxic conditions ^35, 36^. We demonstrate a novel mode of BNIP3 degradation that is lysosome-mediated but autophagy-independent. This pathway requires ER insertion by the ER membrane protein complex (EMC) and subsequent trafficking through the canonical secretory pathway. Endolysosomal regulation works alongside, but independent of, UPS-mediated regulation of BNIP3, providing an additional regulatory axis for governing BNIP3-mediated mitophagy and its associated physiology. In the process, we directly implicate endosomal trafficking and lysosomal degradation in the canon of quality control pathways that ensure proper localization of TA membrane proteins.

## RESULTS

### Lysosomal delivery of BNIP3 is independent of autophagy

Lysosomal degradation of autophagy receptors is a common proxy for selective autophagy. Using this rationale, we set out to monitor lysosomal delivery of endogenous BNIP3. To this end, we used MDA-MB-231 cells, a triple-negative breast cancer cell line that prominently expresses BNIP3. As previously reported, BNIP3 appears as multiple bands via immunoblot, reflective of variably phosphorylated species, which we confirmed by an *in vitro* dephosphorylation assay (Fig S1A)^16, 37^. BNIP3 protein levels accumulated in MDA-MB-231 cells treated with Bafilomycin-A1 (Baf-A1), a V-ATPase inhibitor that blocks lysosomal acidification, confirming that BNIP3 is degraded in a lysosome-dependent manner (Fig 1A). To test if this lysosomal delivery was mediated by autophagy, we transduced Cas9-expressing cells with a single-guide RNA (sgRNA) targeting *ATG9A*, a core autophagy component, and selected in puromycin for 8 days to generate a non-clonal knockout population. Unexpectedly, the deletion of *ATG9A* did not affect BNIP3 protein levels or its response to Baf-A1 treatment. A similar trend was observed for the related mitophagy receptor, NIX. Importantly, canonical selective autophagy receptors p62 and NDP52 accumulated upon either Baf-A1 treatment or sgATG9A transduction as expected for *bona fide* autophagy substrates (Fig 1A). Comparable results were obtained from a clonal *ATG9A^KO^* isolate (Fig. S1B).

**Figure 1:**
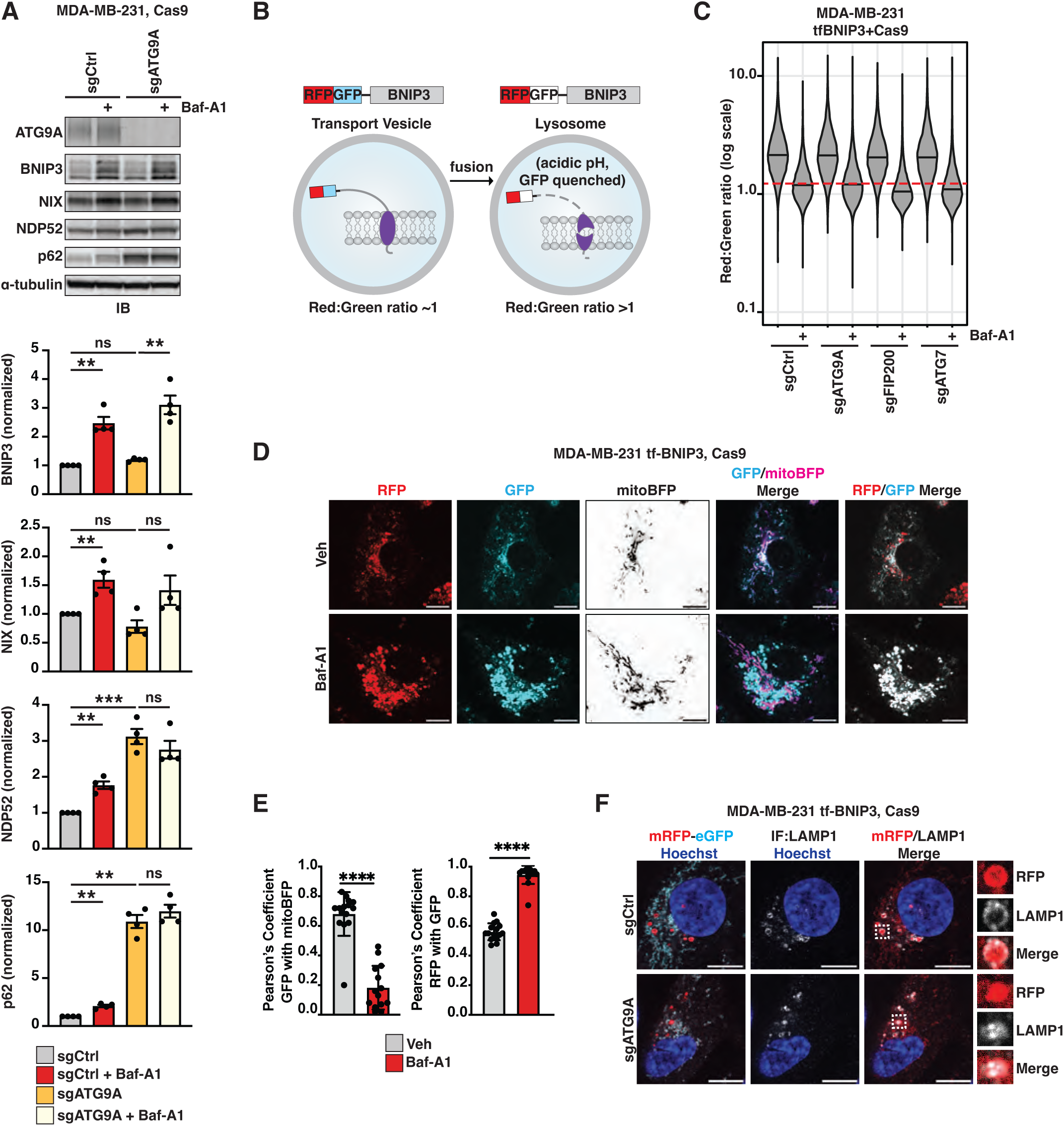
Lysosomal delivery of BNIP3 is independent of autophagy. **(A)** Immunoblotting (IB) of MDA-MB-231-derived extracts from cells expressing Cas9 and the indicated sgRNA. Where indicated, cells were treated with 100nM Baf-A1 for 18hr. Shown are representative images from one biological replicate. Bar graphs represent mean +/- SEM from 4 independent experiments. All protein levels were normalized to α-tubulin. Statistical analysis was performed using a one-sample t-test to the normalized control and an unpaired Student’s t-test between experimental samples. Ctrl, non-targeting control. ***, *p* < 0.001; **, *p* < 0.01; *ns*, not significant. **(B)** Schematic of the tf-BNIP3 reporter. Upon lysosomal delivery, GFP fluorescence is selectively quenched. Thus, corresponding changes in red:green ratio reflect delivery to lysosomes. **(C)** tf- BNIP3-expressing cells were transduced with the indicated sgRNAs. Cells were subsequently treated with DMSO or Baf-A1 (100nM) for 18hr before being analyzed by flow cytometry for red:green ratio. Median values for each sample are identified by a black line within each violin. The red dotted line across all samples corresponds to red:green ratio of maximally inhibited conditions (Baf-A1) (n > 10,000 cells). **(D)** Representative confocal micrographs of tf-BNIP3 cells transiently expressing mitoBFP. Cells were treated with vehicle (DMSO) or Baf-A1 (100nM) for 18hr prior to imaging. Scale bar: 10*µ*m. **(E)** Quantification of Pearson’s correlation coefficients from cells in **D**. Correlation of RFP with GFP (an anti-correlate of lysosomal delivery) and GFP to mitoBFP (reflective of mitochondrial localization) was calculated using Coloc2. Bar graphs represent mean +/- SEM. Each data point represents a single cell. n = 15 cells. Statistical analysis was performed using an unpaired t-test. ****, *p* < 0.0001. **(F)** Representative confocal micrographs of cells transduced with sgRNA constructs targeting *ATG9A* or a non-targeting control (Ctrl). Cells were fixed 8 days post-transduction and immunostained for LAMP1 prior to imaging. Scale bar: 10*µ*m.

Because hypoxia induces BNIP3-and NIX-mediated mitophagy, we reasoned that autophagy-dependent lysosomal delivery of these factors might occur preferentially under hypoxic conditions. To test this, we incubated cells in low oxygen (1% O2) for 18hr, whereupon we observed an increase in BNIP3 protein levels consistent with known transcriptional regulation (Fig S1C). Regardless, *ATG9A* still did not affect BNIP3 protein levels relative to control cells (Fig S1C). This autophagy-independent lysosomal degradation of BNIP3 was observed across a diverse panel of cell lines including U2OS, HEK293T, MDA-MB-435, and K562 (Fig. S1D-F). From this, we conclude that BNIP3 (and to a lesser extent, NIX) constitutively undergo robust lysosomal-mediated degradation that is primarily independent of autophagy.

To better dissect the lysosomal delivery of BNIP3, we adapted a tandem fluorescent (tf) system composed of a red fluorescent protein (RFP) and a green fluorescent protein (GFP) fused to a protein of interest, in this case BNIP3 ^38, 39^. GFP fluorescence is selectively quenched in the low pH environment of the lysosomal lumen. In contrast, RFP fluorescence persists (Fig 1B). Therefore, the red:green ratio serves as a ratiometric proxy for lysosomal delivery and can be quantified with single-cell resolution. Utilizing the tf-reporter system, we generated Cas9-expressing MDA-MB-231 cells stably co-expressing N-terminally tagged BNIP3 from the AAVS1 safe-harbor locus. By this approach, we observed a striking collapse in the red:green ratio of our tf-BNIP3 reporter in cells treated with Baf-A1, consistent with our earlier observations (Fig 1C). In contrast, inhibiting autophagy with a chemical inhibitor of VPS34, PIK-III, failed to collapse the red:green ratio of tf-BNIP3, despite inhibiting flux of a canonical autophagy reporter, tf-NDP52 (Fig S1G)^40^. By a complementary genetic approach, we similarly found that knockdown of Rab7A, a small GTPase broadly associated with the late endosomal system, collapsed the red:green ratio of tf-BNIP3 (Fig. S1H), while tf-BNIP3 flux persisted cells lacking key autophagy-specific factors: ATG9A, FIP200, or ATG7 (Fig 1C). To further validate the tf-BNIP3 reporter, we monitored tf-BNIP3 expression and localization in MDA-MB-231 cells using fluorescence-based confocal microscopy. In control cells, GFP signal strongly correlated with mito-BFP, evidence that tf-BNIP3 localizes appropriately to mitochondria (Fig 1D, Fig 1E)^41^. RFP-only puncta were prevalent in DMSO-treated controls but fully collapsed into RFP^+^/GFP^+^ puncta upon Baf-A1 treatment (Fig 1D-E). These RFP-only puncta co-localized with a lysosomal marker, LAMP1, consistent with the interpretation that RFP-only structures reflect lysosomal delivery of the reporter (Fig 1F). Similar results were observed in *ATG9A^KO^* cells, reinforcing that this process is autophagy-independent. As an aside, we note that Baf-A1 treatment depleted the correlation coefficient of RFP or GFP with mito-BFP, suggesting that lysosomally destined RFP^+^/GFP^+^ structures (i.e., BNIP3) do not contain luminal mitochondrial content (Fig 1E, Fig S1I). Collectively, these data indicate that our tf-BNIP3 reporter recapitulates the autophagy-independent degradation of BNIP3 by the lysosome.

### Genome-wide CRISPR screening reveals modifiers of BNIP3 flux

In the absence of an autophagy-mediated pathway, it was uncertain how an outer mitochondrial membrane (OMM) protein would be robustly degraded by the lysosome. To identify factors required for the lysosomal delivery of BNIP3, we employed our tf-BNIP3 reporter to perform a genome-wide CRISPR knockout screen for modifiers of BNIP3 flux. MDA-MB-231 cells expressing Cas9 and tf-BNIP3 were transduced with a lentiviral library containing 76,441 sgRNAs spanning the entire human genome^42^ (Fig 2A). Cells were then sorted by red:green ratio to collect the top and bottom 30% of cells, representing cells that were enhanced and inhibited for lysosomal delivery of tf-BNIP3, respectively (Fig 2A). To identify genes associated with each effect, sgRNAs from each pool were amplified, sequenced, and analyzed with the Model-based Analysis of Genome-wide CRISPR-Cas9 Knockout (MAGeCK) pipeline^43–45^(Table S1). We utilized fold change as a proxy for the strength of a gene as an effector of lysosomal delivery. A negative fold change indicates the gene mediates lysosomal delivery, as the perturbation leads to decreased flux. A positive fold change indicates genes that, when knocked out, induce flux. We categorized the two populations as potential “effectors” and “suppressors”, respectively.

**Figure 2:**
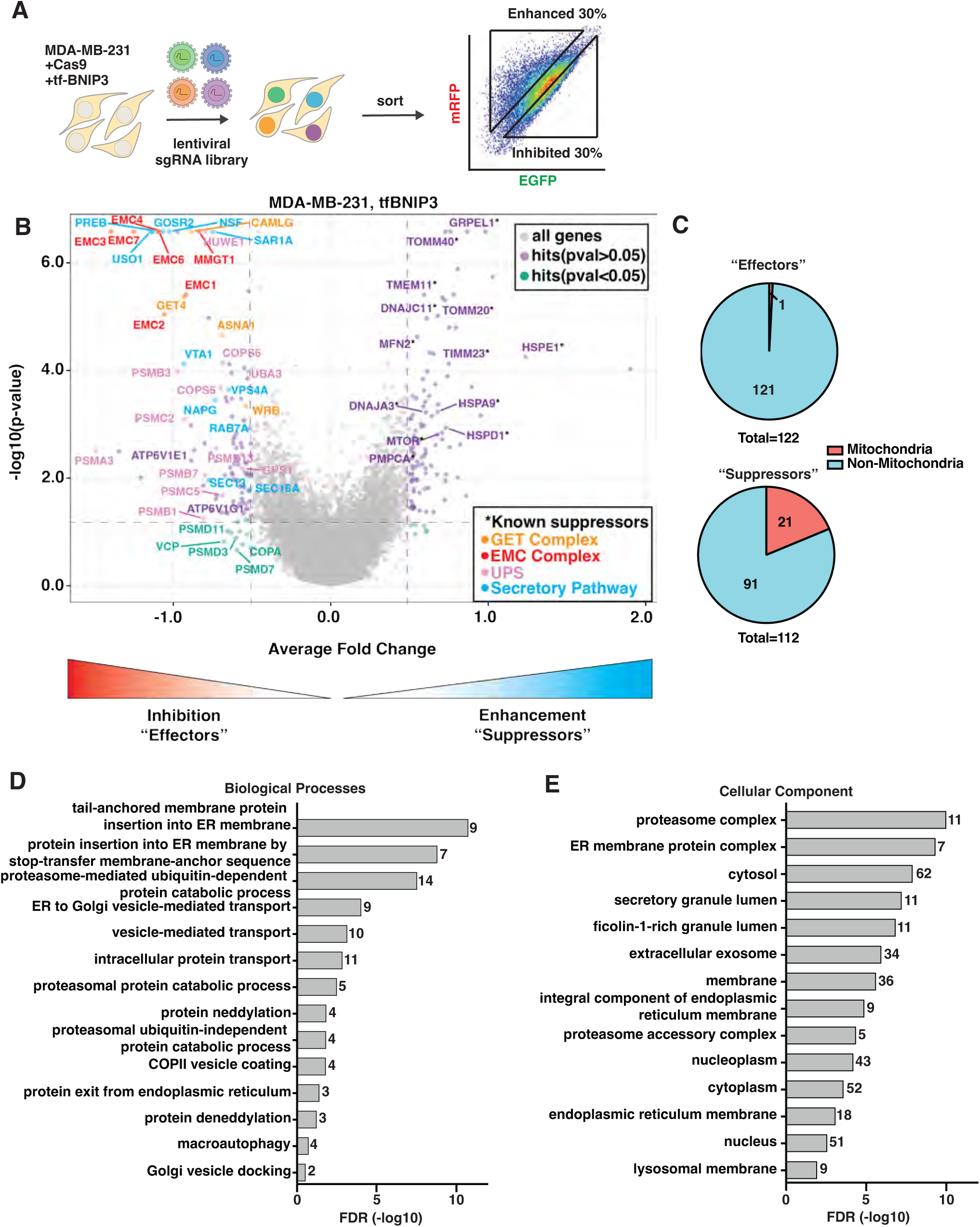
Genome-wide CRISPR screening reveals modifiers of BNIP3 flux. **(A)** Schematic of the genome-wide CRISPR screening pipeline for modifiers of tf-BNIP3 delivery to the lysosome. Reporter cells were transduced with an sgRNA library, propagated, and sorted to collect the top 30% (enhanced delivery) and bottom 30% (inhibited delivery) of tf-BNIP3 cells based on the red:green fluorescence ratio. **(B)** Volcano plot of BNIP3 effectors based on average fold change (a proxy for effect strength) and *p*-value. Average fold changes greater than 0.5 and less than -0.5 are indicated vertical dashed lines. Horizontal dashed line indicates a *p*-value of 0.05. Only genes from cellular pathways or protein complexes validated by this study or independent studies are labeled. **(C)** Effectors and suppressors identified in **B** were mapped against the MitoCarta 3.0 database to identify known mitochondrial factors. **(D-E)** Unbiased Gene Ontology (GO) term analysis of genes in the effector population.

Any gene with a fold change less than -0.5 or greater than 0.5 was considered a “hit” in the screen. At this threshold, we identified 122 effector genes and 112 suppressor genes (Fig 2B, Table S1). Concordant with our preliminary observations, core autophagy factors were absent from the effector population. Yet we recovered Rab7A as an effector, as previously validated (Fig S1H). In addition, multiple suppressor genes identified from our screen had previously been reported including TMEM11, DNAJA3, DNAJC11, and HSPA9^46, 47^. In all, our list of identified effector and suppressor proteins was largely concordant with available data, validating our approach.

Surprisingly, when compared to the MitoCarta 3.0 database^48^, only 1 of 122 effector genes and 21 of 112 suppressor genes were annotated as mitochondrial (Fig 2C). To identify other pathways or components implicated by our data, we performed an unbiased Gene Ontology (GO) analysis. Enriched GO terms in the effector population related to membrane insertion, vesicle-mediated transport, and proteasomal pathways, with many terms specifically pertaining to the endoplasmic reticulum (ER) (Fig 2D, Fig 2E). Previously profiled autophagy receptors do not similarly enrich for these GO terms, suggesting a uniqueness to BNIP3^38^. In particular, our data identified the ER membrane protein complex (EMC), the guided entry of tail-anchored proteins (GET) complex, ER-Golgi transport, and the ubiquitin-proteasome system (UPS) as potential effectors of BNIP3 stability (Fig 2B). In sum, our genetic screening approach identified numerous known regulators of BNIP3 as well as a unique role for ER insertion and ER-to-Golgi trafficking in BNIP3 regulation.

### Lysosomal delivery of BNIP3 is governed by the EMC and the secretory pathway

To validate our screen results, we transduced our tf-BNIP3 reporter cells with a representative subset of individual sgRNAs and monitored corresponding changes in red:green ratio using flow cytometry. These data clearly verified the EMC as a potent effector of BNIP3 degradation as the deletion of EMC subunits mirrors the effect of Baf-A1 treatment (Fig 3A, Fig S2A). In addition, knockout of the GET complex, components of the secretory pathway including multivesicular bodies (MVBs), UPS factors, and vacuolar ATPase subunits all decreased red:green ratio (Fig 3A, Fig S2A-B). Similar effects were observed in U2OS osteosarcoma cells expressing tf-BNIP3 and Cas9, confirming that the effectors we identified are not strictly cell-type specific (Fig S2C).

**Figure 3:**
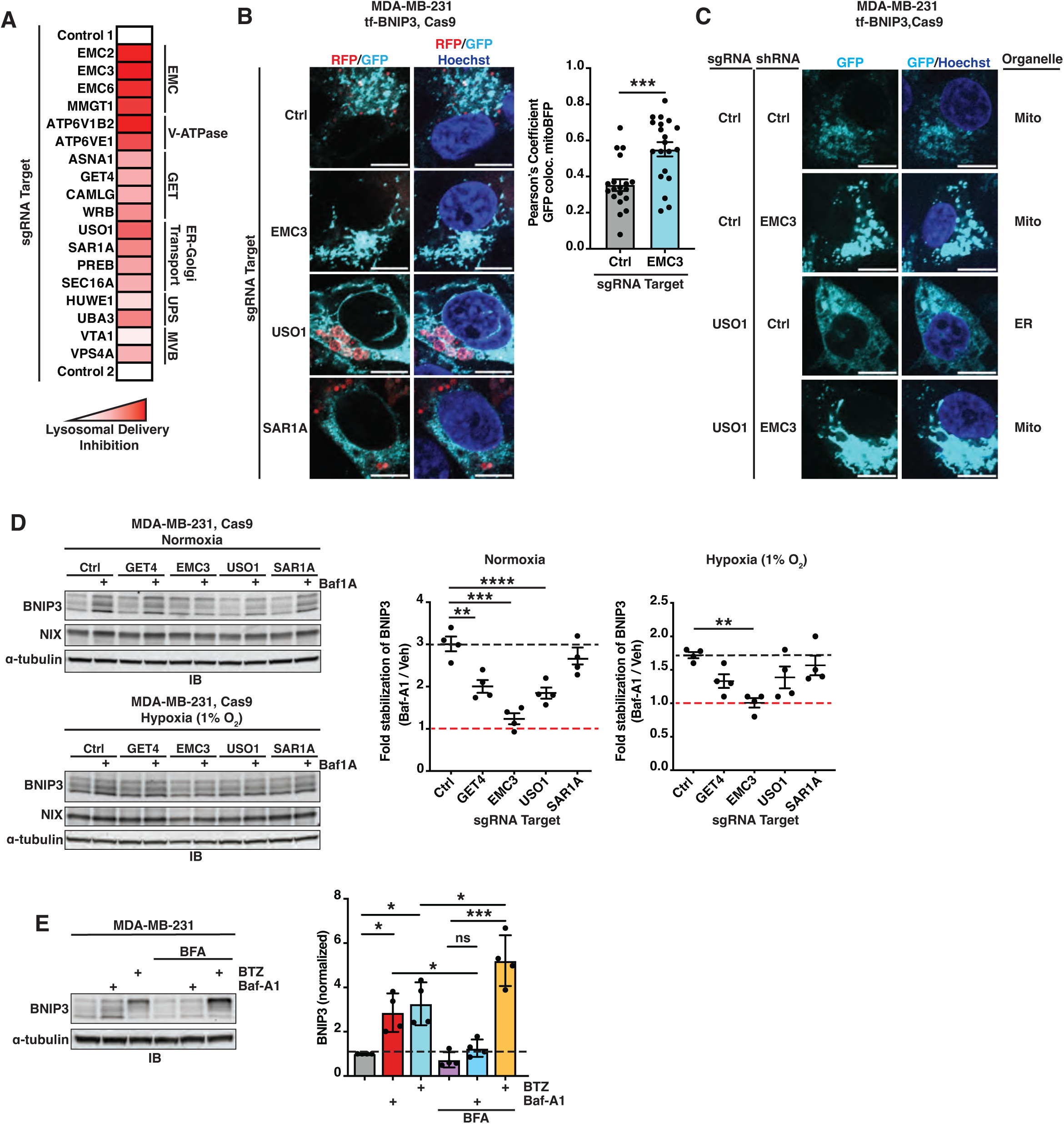
BNIP3 lysosomal delivery is governed by ER-insertion and the secretory pathway. **(A)** MDA-MB-231 cells expressing tf-BNIP3 and Cas9 were transduced with sgRNAs for the indicated genes. The median red:green ratio of each population was used to generate a heatmap. Darker shades of red indicate greater inhibition, with a red:green ratio of 1 taken as the theoretical maximum inhibition. Genes were clustered by related function. For underlying data, see Fig S2A**. (B)** Representative confocal micrographs of tf-BNIP3-expressing cells transduced with sgRNAs targeting the indicated genes. Pearson’s correlation coefficient between GFP and mitoBFP (reflective of mitochondrial localization) was calculated using Coloc2. Bar graphs represent mean +/- SEM. Each data point represents a single cell. Statistical analysis was performed using an unpaired t-test. Scale bar: 10*µ*m; *n* = 15 cells; ***, *p* < 0.001. **(C)** Representative confocal micrographs of tf-BNIP3-expressing cells transduced with indicated sgRNA and shRNA. Scale bar: 10*µ*m **(D)** Immunoblotting (IB) of MDA-MB-231-derived extracts from cells transduced with indicated sgRNAs in both normoxia and hypoxia. Where indicated, cells were treated with 100nM Baf-A1 for 18hr. Shown are representative images from one biological replicate. Quantification of BNIP3 protein stabilization by Baf-A1 treatment was calculated as: (BNIP3^Baf-A^^1^/tubulin^Baf-A^^1^)/(BNIP3^DMSO^/tubulin^DMSO^). Graphs represent the mean +/- SEM from 4 independent experiments. Black dashed line indicates fold-stabilization of BNIP3 upon Baf-A1 treatment in control cells. Red dashed line demarcates no stabilization. Statistical analysis was performed a one-way ANOVA with Tukey’s test. ****, *p* < 0.0001; ***, *p* < 0.001; **, *p* < 0.01. **(E)** Immunoblotting (IB) of MDA-MB-231-derived extracts from cells treated with Brefeldin-A (BFA) (1*µ*M), Baf-A1 (100nM), or bortezomib (BTZ) (100nM) for 18hr. Shown are representative images from one biological replicate. Bar graphs represent mean +/- SEM from 4 independent experiments. All protein levels were normalized to α-tubulin. Statistical analysis was performed using a one-sample t-test to the normalized control and an unpaired Student’s t-test between experimental samples, Veh (DMSO) ***, *p* < 0.001; **, *p* < 0.01; *, *p* < 0.05.

Within the endolysosomal system implicated above, the EMC and GET complex are related ER insertion pathways for tail-anchored proteins^26, 27, 49–51^. Notably, BNIP3 was previously observed on the ER membrane and accumulates on the ER during stress conditions^7, 37, 52, 53^. In *EMC3^KO^*cells, tf-BNIP3 displayed a striking decline in RFP-only puncta with a concomitant increase in the co-localization of BNIP3 with mito-BFP (Fig 3B). Knockout of vesicular transport factors *USO1* or *SAR1A* failed to fully prevent lysosomal delivery of BNIP3. However, both exhibited a marked shift in BNIP3 localization to structures resembling the ER network (Fig 3B). Taking advantage of these differences in localization, we performed an epistasis analysis through pairwise depletion of *EMC3* and *USO1*. While knockout of *USO1* shifts the distribution of BNIP3 primarily to an ER-like morphology, when combined with the knockdown of *EMC3*, BNIP3 shifted back to a primarily mitochondrial localization (Fig 3C). This epistatic relationship suggests the EMC governs BNIP3 entry into the ER membrane, which precedes the role of USO1 and the secretory pathway in trafficking BNIP3 to the lysosome.

To test whether BNIP3 trafficking through the endolysosomal system was an artifact of over-expression, we transduced Cas9-expressing MDA-MB-231 cells with sgRNAs targeting GET4, EMC3, USO1, or SAR1A and selected under puromycin for 8 days. We then subjected these cells to normoxic or hypoxic conditions and monitored cellular extracts for changes in endogenous BNIP3 levels. All responses were measured in comparison to chemical inhibition of the lysosome by Baf-A1. When comparing each Baf-A1-treated knockout to its respective DMSO-treated control, we saw Baf-A1 sensitivity diminish (Fig 3D). Knockout of *EMC3* remained the most potent effector, as BNIP3 protein levels were completely insensitive to Baf-A1 treatment in this background. Knockout of *GET4* and *USO1* resulted in a reduced sensitivity to Baf-A1, while knockout of *SAR1A* had only a minimal effect (Fig 3D). Similar trends were observed in U2OS cells (Fig S2D). These results affirm that deletion of the EMC prevents lysosomal delivery of BNIP3.

As an independent measure of the role of the secretory pathway in delivering BNIP3 to lysosomes, we utilized a chemical inhibitor of ER-to-Golgi transport, Brefeldin-A (BFA). Treatment with BFA alone had no significant effect on endogenous levels of BNIP3. However, BFA treatment fully negated the stabilizing effects of Baf-A1, consistent with the model that ER-to-Golgi trafficking of BNIP3 is a prerequisite for its lysosomal delivery. In contrast, BFA treatment potentiated the effect of bortezomib (BTZ), a proteasome inhibitor, on BNIP3 accumulation (Fig 3E). Collectively, these results reveal that when endolysosomal transport of BNIP3 is perturbed, BNIP3 can no longer be degraded by the lysosome, although it is re-routed for proteasomal degradation.

### Proteasomal degradation restricts BNIP3 levels but not lysosomal delivery

Post-translational control of BNIP3 stability was previously reported to depend on the ubiquitin-proteasome system^12–16, 54–56^. Consistent with these reports, our list of genetic effectors recovered numerous UPS factors previously implicated in the regulation of BNIP3, including proteasomal subunits, the NEDD8 conjugation machinery, and the membrane protein extratase valosin-containing protein (VCP). Indeed, we found that BNIP3 protein levels accumulated upon chemical inhibition of either neddylation (MLN4924) or VCP (CB-5083), particularly under hypoxic conditions (Fig 4A, S3A-B). Thus, our data support the recently emerging role of the UPS in broadly regulating BNIP3.

**Figure 4:**
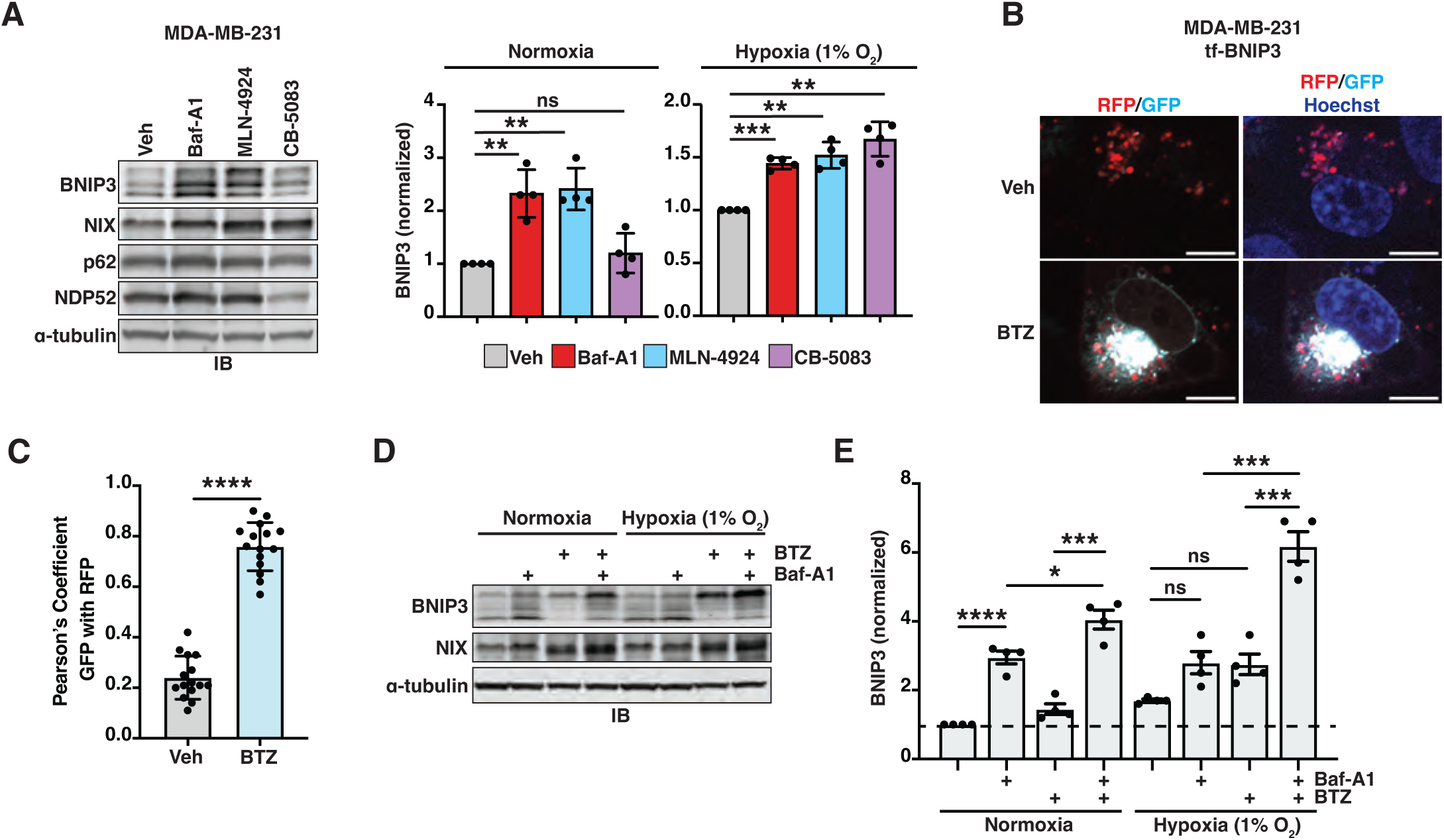
The proteasome is required for efficient BNIP3 degradation, but not lysosomal delivery. **(A)** Immunoblotting (IB) of MDA-MB-231-derived extracts from cells treated with vehicle (DMSO), Baf-A1 (100nM), MLN-4924 (1*µ*M), and CB-5083 (1*µ*M) for 18hr. Shown are representative images from one biological replicate (for hypoxia, see Fig S3A). Bar graphs represent mean +/- SEM from 4 independent experiments. All protein levels were normalized to α-tubulin. Statistical analysis was performed using a one-sample t-test to the normalized control and an unpaired Student’s t-test between experimental samples test. ***, *p* < 0.001, **; *p* < 0.01; *ns*, not significant. **(B)** Representative confocal micrographs of fixed tf-BNIP3-expressing cells treated with vehicle (DMSO) or bortezomib (BTZ) (100nM) for 18hr. Scale bar: 10*µ*m. **(C)** Pearson’s correlation coefficient between GFP and RFP was calculated using Coloc2. Bar graph represents mean +/- SEM. Each data point represents a single cell. Statistical analysis was performed using an unpaired Student’s t-test. Scale bar: 10*µ*m; *n* = 15 cells; ****, *p* < 0.0001. **(D)** Immunoblotting (IB) of MDA-MB-231-derived extracts from cells grown in normoxia and hypoxia and treated with DMSO, Baf-A1 (100nM), and/or bortezomib (BTZ) (100nM) for 18hr. Shown are representative images from one biological replicate. **(E)** Quantification of BNIP3 protein accumulation from **D**. Bar graphs represent mean +/- SEM from 4 independent experiments. All protein levels were normalized to α-tubulin. Statistical analysis was performed using a one-way ANOVA with Dunnett’ test and an unpaired Student’s t-test between experimental samples. ****, *p* < 0.0001; ***, *p* < 0.001; *, *p* < 0.05; *ns*, not significant.

Based on the data above, we wished to better explore the interplay between proteasomal and lysosomal regulation of BNIP3. Using fluorescence microscopy, we saw a striking stabilization of tf-BNIP3 intensity upon proteasomal inhibition, but we still noted the presence of RFP-only puncta (Fig 4B-C). We interpret this to indicate that proteasomal inhibition dramatically stabilizes tf-BNIP3 protein levels, but it does not arrest lysosomal delivery *per se*. To test this, we grew parental MDA-MB-231 cells in normoxic or hypoxic conditions, with or without Baf-A1 and/or BTZ for 18hr. We then monitored extracts by immunoblotting for endogenous levels of BNIP3 and NIX (Fig 4D). As reported above (Fig 1D), Baf-A1 significantly stabilized BNIP3 levels regardless of oxygen availability (Fig 4E). Likewise, BTZ had a generally stabilizing effect that was comparable to or lesser than Baf-A1. We note that the qualitative changes in the BNIP3 banding pattern reflect a hyper-phosphorylated species that appears upon proteasome inhibition (Fig 4D, Fig S1A). When combined, treatment with Baf-A1 and BTZ resulted in the additive accumulation of BNIP3 under both normoxic and hypoxic conditions, supporting non-overlapping roles for the proteasome and the endolysosomal system in restricting BNIP3 levels (Fig 4E).

### BNIP3 dimerization determines its mode of degradation and is required for lysosomal delivery

The soluble portion of BNIP3 (residues 1-163) contains several known motifs and domains. BNIP3 contains a canonical LC3-interacting region (LIR) motif required for mitophagy^6^. In addition, it contains a PEST domain, a BH3 domain, and an extended “conserved region” adjacent to the BH3 domain (Fig 5A)^18, 57^. To elucidate the features within BNIP3 that are required for its lysosomal delivery, we performed a structure-function analysis using our tf-reporter system (Fig 5A). To this end, we transient expressed tf-BNIP3 variants and monitored red:green ratio as a proxy for lysosomal delivery. Mutation of the LIR motif (W18A/L21A) or truncation of the LIR motif (aa30-end) had little effect on the lysosomal delivery of BNIP3. Similarly, a phospho-mimetic mutation near the LIR motif, BNIP3^S17E^, that enhances LC3 binding did not increase lysosomal delivery (Fig 5B)^6^. Additional truncations of the PEST domain (aa82-end or aa104-end) also had minimal effect on flux. Subsequent deletion of the BH3 domain (aa117-end) partially diminished lysosomal delivery although delivery was still active (Fig 5B). The BH3 domain also has been implicated in the proteasomal regulation of the BNIP3, which we confirmed (Fig S4A)^16^. Only a near-complete truncation of the soluble domain (aa137-end), which also eliminates the conserved region, showed dramatic inhibition of lysosomal delivery. Concordant with its decrease in lysosomal delivery, the aa137-end truncation exhibited an increasingly mixed mitochondrial/ER localization pattern, including significant signal on the perinuclear membrane (Fig S4B). We conclude that both the conserved region and, to a lesser extent, the BH3 domain, influence the lysosomal trafficking of BNIP3, likely by facilitating ER export.

**Figure 5:**
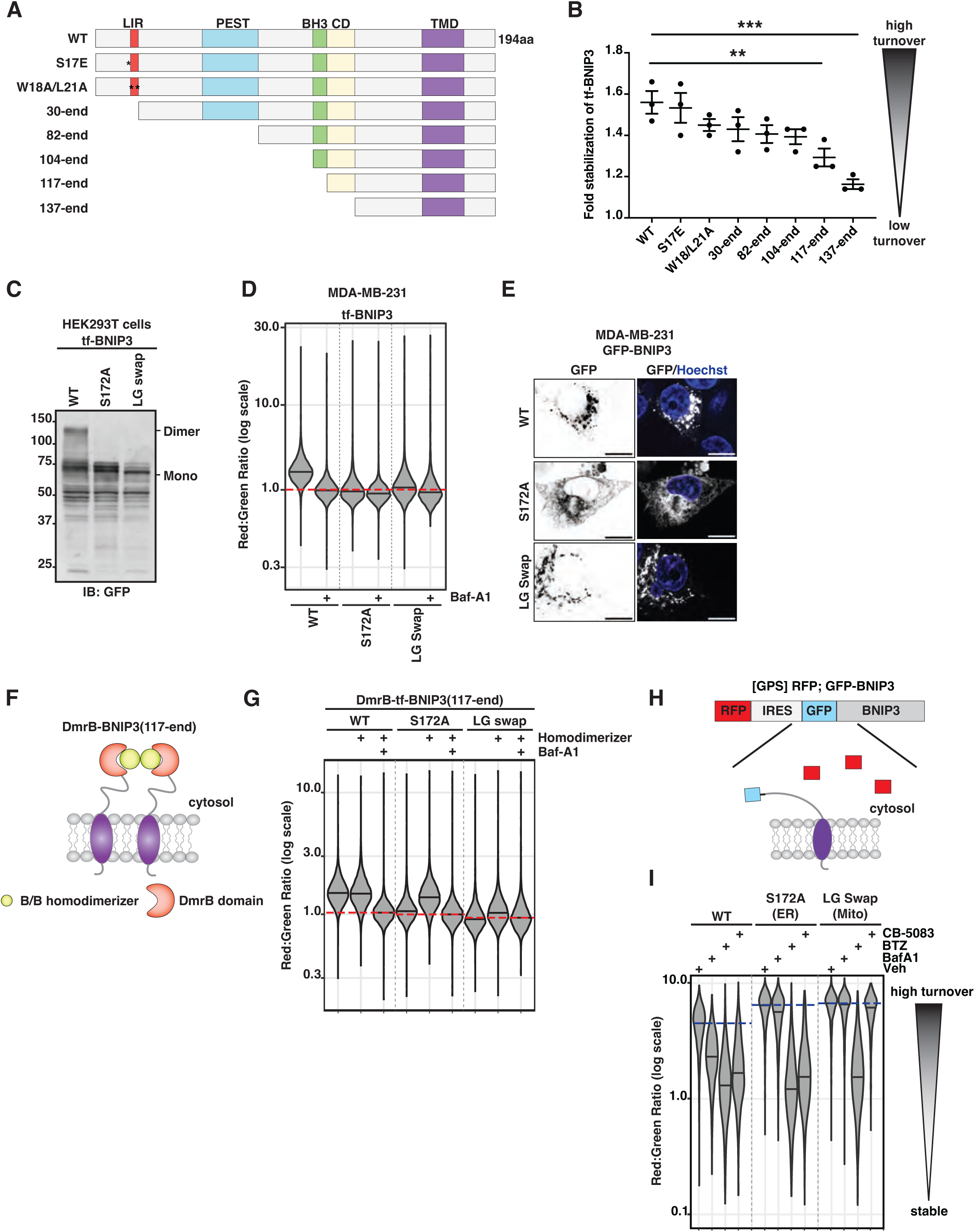
BNIP3 dimerization determines mode of degradation and is required for lysosomal delivery. **(A)** Domain organization of BNIP3 (NP_004043.4) and derived variants. LC3, LC3-interacting region; PEST, PEST domain; BH3, BH3 domain; CR, conserved region; TMD, transmembrane. **(B)** Dot plot representing fold-stabilization of tf-BNIP3 variants in response to Baf-A1. Stabilization was calculated as a ratio of median red:green ratios (DMSO/Baf-A1). A ratio of 1 represents no lysosomal delivery. Statistical analysis was performed using a one-way ANOVA with Dunnett’ test. ***, *p* < 0.001; **, *p* < 0.01. **(C)** Immunoblotting (IB) of HEK293T-derived extracts transiently expressing the indicated tf-BNIP3 variants. Monomeric and dimeric species are indicated. **(D)** MDA-MB-231 cells were transduced with the indicated tf-BNIP3 variants. Red:green ratio was analyzed by flow cytometry 48hr post-transduction. Cells were incubated with vehicle (DMSO) or Baf-A1 (100nM) for 18hr before analysis. Median values for each sample are identified by a black line within each violin. The red dotted line across all samples corresponds to red:green ratio of wild-type (WT) cells inhibited with Baf-A1 (n > 10,000 cells). **(E)** Representative confocal micrographs of MDA-MB-231 cells transduced with indicated GFP-BNIP3 variants. Scale bar, 10*µ*m. **(F)** Schematic of the DmrB-based inducible dimerization system using the 117-end variant of BNIP3. **(G)** MDA-MB-231 cells were transduced with the indicated tf-BNIP3^117^–^end^ variants. Red:green ratio was analyzed by flow cytometry 48hr post-transduction. Cells were treated with Baf-A1 (100nM) and/or B/B homodimerizer (0.5 *µ*M) for 6hr prior to performing flow cytometry. Median values for each sample are identified by a black line within each violin. The red dotted line across all samples corresponds to wild-type (WT) cells inhibited with Baf-A1 (n > 10,000 cells). **(H)** Schematic of GPS cassette fused to BNIP3. IRES, internal ribosome entry site. **(I)** MDA-MB-231 cells were transduced with the indicated [GPS]BNIP3 variants. Red:green ratio was analyzed by flow cytometry 48hr post-transduction. Cells were treated with vehicle (DMSO), Baf-A1 (100nM), BTZ (100nM), and CB-5083 (1*µ*M) for 18hr prior to performing flow cytometry. Median values for each sample are identified by a black line within each violin. The red dotted line across each sample group corresponds to the basal red:green ratio of mock-treated cells (n > 10,000 cells).

As a representative tail-anchored protein, BNIP3 also possesses a single, C-terminal, transmembrane domain that is essential for its localization, insertion into membranes, and dimerization^5, 18, 58^. As dimerization has been routinely tied to the functionality of both BNIP3 and NIX^59–61^, we generated two transmembrane mutants in BNIP3 to investigate the role of dimerization in lysosomal delivery. First, we generated a frequently used serine-to-alanine mutation (S172A), which disrupts intermonomer side chain hydrogen bonding^62–64^. Second, we swapped the positions of leucine-179 and glycine-180 (LG swap). These two residues are part of the transmembrane GxxxG motif required for dimer formation^18, 63, 64^. The LG swap mutation disrupts the motif registrar while maintaining the overall hydrophobicity of the TMD segment. When expressed in HEK293T cells, both mutations disrupted the formation of SDS-resistant dimers (Fig 5C). In a corresponding functional assay, both dimer mutations disrupted BNIP3 delivery to lysosomes (Fig 5D). We then monitored the cellular localization of the two dimerization mutants to see where they arrested. Surprisingly, our dimerization mutants were differentially localized. BNIP3^S172A^ localized in a reticular, ER-like pattern (Fig 5E). We anticipate this shift in localization is due to the changing hydrophobicity of the transmembrane domain, as hydrophobicity is a primary determinant for tail-anchored protein targeting^22^. In contrast, the LG swap of the GxxxG motif, which does not affect hydrophobicity, remained primarily on mitochondria (Fig 5E).

Failure of dimerization mutants to traffic to the lysosome suggests that dimerization is an important aspect of BNIP3 trafficking. However, localization discrepancies limited our ability to cleanly interpret these results. To solidify the role of dimerization in BNIP3 trafficking, we turned to an inducible dimerization system (Fig 5F)^65^. In short, a DmrB domain was fused onto the N-terminus of the shortest functional BNIP3 truncation (aa117-end). This allowed us to position the artificial dimerization domain as close to the transmembrane helix as possible. Next, we transiently expressed tf-BNIP3^117^–^end^ or the dimerization mutants (S172A and LG swap) with or without an in-frame DmrB dimerization domain. We then incubated cells with a small molecule homodimerizer and monitored red:green ratio as a proxy for flux. The homodimerizer molecule did not affect the red:green ratio for DmrB-tf-BNIP3^117^–^end^ or any constructs lacking the DmrB domain (Fig 5G, S4C). However, incubation with homodimerizer rescued the red:green ratio of the DmrB-fused S172A mutant to that of the wild type (Fig 5G). Importantly, Baf-A1 attenuated this increase, confirming the increase was due to lysosomal delivery. In contrast, the mitochondrially-restricted LG swap mutant was minimally responsive to the homodimerizer (Fig 5G). Collectively, these data illustrate that dimerization within the ER membrane is a required aspect of BNIP3 trafficking to the lysosome.

Within this model, what is the fate of unassembled BNIP3 monomers? To address this, we employed the global protein stability (GPS) cassette, a reporter used to study proteasomal degradation and protein degrons^66^. In brief, the cassette contains an RFP fluorophore, followed by an internal ribosome entry site (IRES) and a GFP fluorophore tethered to BNIP3 (Fig. 5H). This results in the expression of two polypeptides: a cytosolic RFP and a GFP-BNIP3 fusion protein. The relative stability of individual GFP-BNIP3 variants can then be assessed by red:green ratio. This approach is methodologically similar to the tf-BNIP3 reporter. However, the output better incorporates the effects of proteosomal regulation. By this approach, we observe that dimerization mutants are destabilized compared to wild type (Fig 5I). Consistent with our tf-BNIP3 reporter, treatment with Baf-A1 stabilized wild-type BNIP3 but did not stabilize either dimer mutant. However, both dimer mutants were dramatically stabilized by a chemical inhibitor of the proteasome, BTZ, indicating they are selectively targeted by the UPS. Chemical inhibition of VCP (CB-5083) selectively stabilized ER-localized monomers, highlighting that proteostatic regulation of BNIP3 is governed by organelle-specific mechanisms (Fig 5I). Collectively, these results suggest that BNIP3 dimerization state and organelle localization determines the mode of degradation.

### Lysosomal delivery of BNIP3 is distinct from BNIP3-mediated mitophagy

Autophagy receptors are frequently degraded in tandem with their cargo. The observation that BNIP3 flux is largely autophagy-independent opposes this paradigm, leading us to more specifically evaluate the relationship between BNIP3 flux and BNIP3-mediated mitophagy. To distinguish the lysosomal delivery of BNIP3 from BNIP3-mediated mitophagy, we turned to an established mitophagy reporter, mt-Keima^67^. This reporter encodes a cytochrome c oxidase signal sequence fused to a pH-sensitive fluorescent protein, mKeima. In MDA-MB-231 cells, we observed moderate basal flux of mt-Keima (15%), as normalized to Baf-A1 treatment (Fig 6A). Knockout of *ATG9A* did not inhibit lysosomal delivery of the mt-Keima reporter (Fig S5A, compare sgCtl vs sgATG9A). Thus, basal flux in MDA-MB-231 cells is largely independent of autophagy. Autophagy-independent delivery of mitochondrial content to lysosomes is likely due to mitochondrial-derived vesicles^68–70^. Therefore, we dubbed the readout for the mt-Keima reporter as “mitoflux” to incorporate autophagic and non-autophagic turnover of mitochondria.

**Figure 6:**
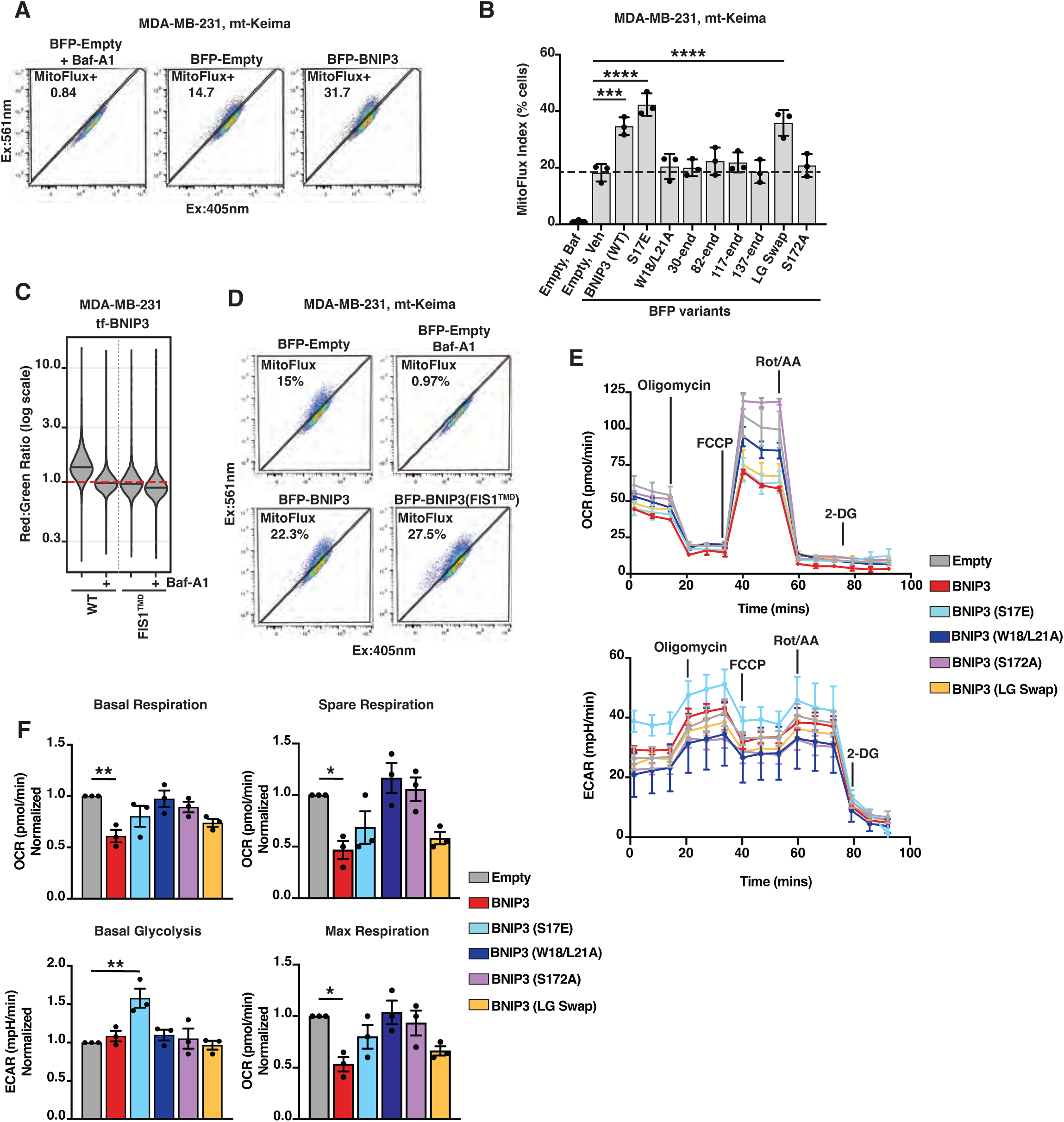
Lysosomal delivery is distinct from BNIP3-mediated mitophagy. **(A)** MDA-MB-231 cells expressing mt-Keima were transduced with BFP-empty or BFP-BNIP3 and analyzed by flow cytometry 48hr post-transduction. Cells were incubated with vehicle (DMSO) or Baf-A1 (100nM) for 18hr before analysis. Baf-A1 treatment was used to define “MitoFlux+”, indicative of cells turning over mitochondria. n > 10,000 cells. **(B)** MDA-MB-231 cells expressing mt-Keima were transduced with indicated BFP-BNIP3 variants and analyzed for Mitoflux as in **A**. Bar graphs represent mean +/-SEM from 3 independent experiments. Statistical analysis was performed using a one-way ANOVA with Dunnett’s test. ****, *p* < 0.0001; ***, *p* < 0.001. **(C)** MDA-MB-231 cells were transduced with wild-type (WT) tf-BNIP3 or tf-BNIP3(FIS1^TMD^) and analyzed by flow cytometry 48hr post-transduction. Cells were incubated with vehicle (DMSO) or Baf-A1 (100nM) for 6hr before analysis. Median values for each sample are identified by a black line within each violin. The red dotted line across all samples corresponds to red:green ratio of wild-type cells inhibited with Baf-A1 (n > 10,000 cells). **(D)** MDA-MB-231 cells expressing mt-Keima were transduced with indicated BFP-BNIP3 variants and analyzed for Mitoflux as in **A**. **(E)** MDA-MB-231 cells were transduced with indicated the BFP-BNIP3 variants and analyzed for oxygen consumption rate (OCR) and extracellular acidification rate (ECAR) 48hr post-transduction. Values were normalized by BCA protein assay. **(F)** Quantification of basal respiration, basal glycolysis, spare respiration, and max respiration from **E**. Bar graphs represent mean +/- SEM from 3 independent experiments. Statistical analysis was performed using a one-way ANOVA with Dunnett’s test. **, *p* < 0.01; *, *p* < 0.05.

To assess how BNIP3 variants influence mitophagy, we took advantage of the fact that BNIP3 overexpression induces mitophagy^9–11^. We transiently expressed BFP-BNIP3 variants or a BFP empty vector control in MDA-MB-231 cells expressing mt-Keima. Expression of wild-type BNIP3 notably increased the percentage of mitoflux+ cells as compared to the BFP-only control (31.7% vs 14.7%) (Fig 6A). Combination treatment with hypoxia led to an additive induction of mitoflux (Fig S5B). We then tested mitophagy induction by BNIP3 mutants. Consistent with previous reports^6, 16^, the phosphomimetic S17E mutant modestly increased mitoflux above wild-type BNIP3 (mean values 18.3% empty vs 34.7% WT vs 42% S17E, p<0.0001) (Fig 6B). Correspondingly, the double LIR mutant (W18/L21A) failed to enhance mitoflux as did all tested truncations of BNIP3 (Fig 6B). We note that these data strongly contrast with the trends observed for endolysosomal trafficking of BNIP3 (compare Fig 6B and Fig 5B), confirming that lysosomal delivery and mitophagy are functionally separable features of BNIP3.

In addition to the LIR motif, the transmembrane helix has been intermittently implicated in BNIP3-and NIX-mediated mitophagy^7, 60^. We found the mitochondrially localized LG swap mutant induced mitoflux comparably to wild type while the ER-localized S172A mutant did not (Fig 6B). This discrepancy suggests that BNIP3-induced mitophagy is a function of localization, not dimerization. To test this, we swapped the endogenous BNIP3 transmembrane domain with the transmembrane domain from an unrelated mitochondrial TA protein, Fis1 (hereafter BNIP3(FIS1^TMD^)). A tf-BNIP3(FIS1^TMD^) chimera failed to traffic to the lysosome, presumably due to abolished dimerization and/or diminished ER localization (Fig 6C). However, BNIP3(FIS1^TMD^) induced mitophagy comparable to wild-type BNIP3 (Fig 6D)^7^. These data indicate that the cytosolic portion of the BNIP3 protein tethered to the OMM is sufficient to induce mitophagy, and the native BNIP3 TMD domain is not required for mitophagy.

Are the aforementioned, BNIP3-dependent, changes in mitoflux sufficient to affect cellular physiology? To evaluate the functional consequences of this mitophagy, we analyzed metabolic flux in cells expressing BNIP3 or its variants. Ectopic expression of BNIP3 variants decreased oxygen consumption rates (OCR) and increased extracellular acidification rate (ECAR) commensurate with their ability to induce mitophagy (Fig 6E-F, S5C). Thus, the levels of mitophagy induced by ectopic BNIP3 expression are sufficient to drive changes in global energy metabolism.

### The endolysosomal and proteosomal systems confine BNIP3 levels to suppress basal mitophagy

While BNIP3-induced mitophagy affects cellular physiology, these results were obtained through ectopic expression of BNIP3. What contribution does endogenous BNIP3 make towards mitoflux and cellular metabolism, and how does regulation by the UPS and the endolysosomal system impinge upon this system? To begin, we induced broad proteostatic collapse in mt-Keima cells using bortezomib, CB-5083, or MLN-4924. Mitoflux increased upon treatment with all three inhibitors (Fig 7A). Moreover, the induction of mitoflux by proteostatic collapse was dependent on *ATG9A* and *BNIP3*, consistent with a mitophagy-specific defect (Fig 7B, S6A). Mitoflux induction by hypoxia displayed similar dependence on *ATG9A* and *BNIP3* (Fig S6B). We then interrogated the role of the endolysosomal system in regulating mitoflux. To this end, we transduced Cas9-expressing mt-Keima cells with an sgRNA targeting *EMC3*. Knockout of *EMC3* induced mitoflux relative to a control sgRNA (∼17% vs ∼34%, p<0.05) and combining *EMC3* deletion with proteasome inhibition had additive effects on mitoflux (30% vs 47.8%, p<0.001)(Fig 7C, S6C). Critically, while *EMC3* deletion elevated mitoflux, this effect was strongly dependent on BNIP3, as concurrent knockdown of BNIP3 with a short hairpin RNA (shBNIP3) returned mitoflux values to baseline (Fig 7D). In sum, these data are consistent with the UPS and the endolysosomal system making non-overlapping contributions towards restricting endogenous BNIP3 mitoflux, and they establish BNIP3 as a node of integration for endolysosomal and proteasomal regulation of mitophagy (Fig 7E).

**Figure 7:**
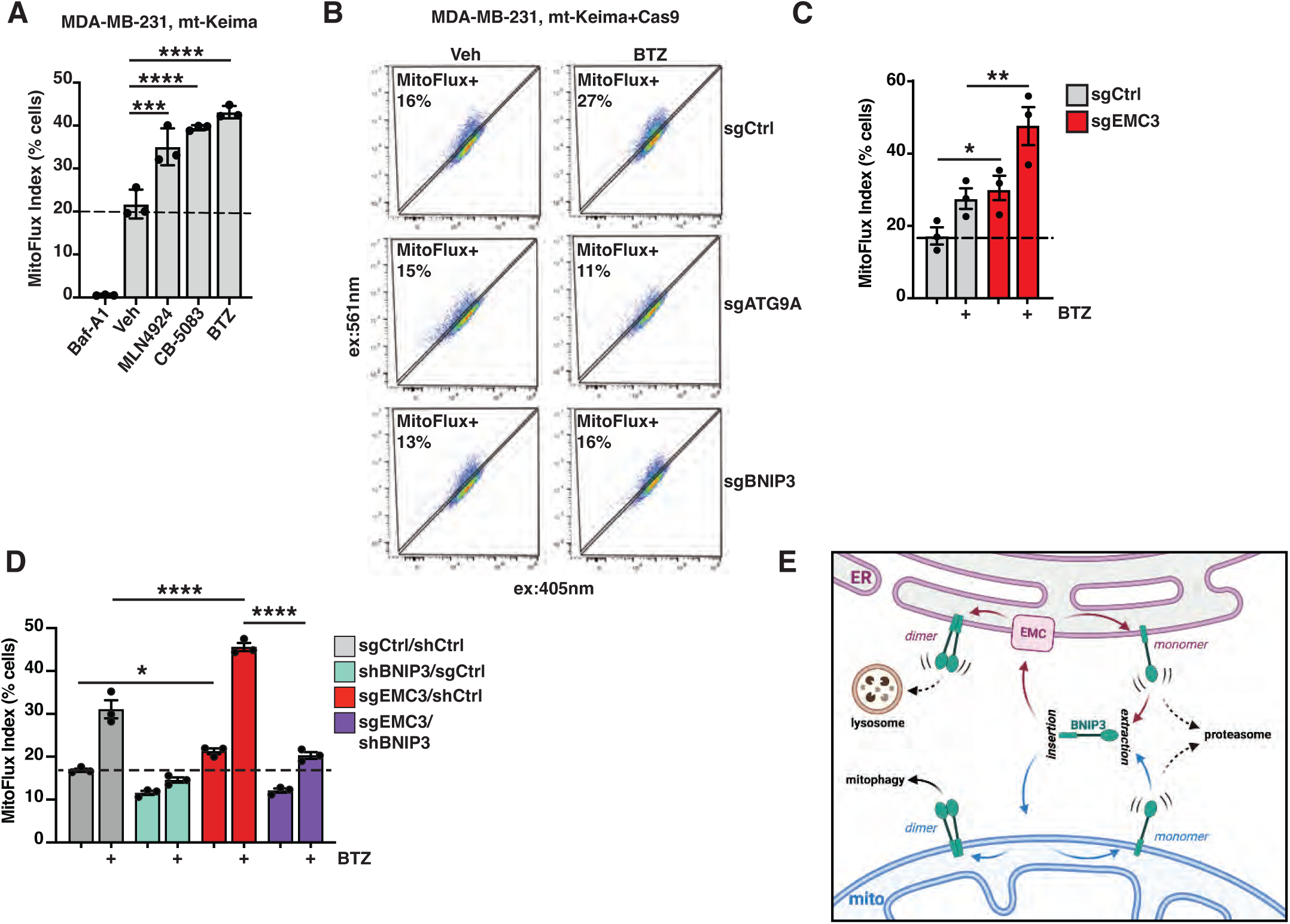
Endolysosomal and proteasomal systems confine BNIP3 levels to suppress basal mitophagy. **(A)** MDA-MB-231 mt-Keima cells treated with vehicle (DMSO), Baf-A1 (100nM), MLN-4924 (1*µ*M), and CB-5083 (1*µ*M) for 24hr prior to analysis by flow cytometry. Bar graphs represent mean +/- SEM from 3 independent experiments. Statistical analysis was performed using a one-way ANOVA with Dunnett’s test. ****, *p* < 0.0001; ***, *p* < 0.001. **(B)** MDA-MB-231 cells expressing mt-Keima were transduced the indicated sgRNAs. Cells were incubated with vehicle (DMSO) or Baf-A1 (100nM) for 18h prior to analysis by flow cytometry. N > 10,000 cells. **(C)** MDA-MB-231 cells expressing mt-Keima were transduced the indicated sgRNAs. Cells were incubated with vehicle (DMSO) or BTZ (100nM) for 18hr prior to flow cytometry. Bar graphs represent mean +/- SEM from 3 independent experiments. Statistical analysis was performed using two-way ANOVA with Tukey’s post-test. **, *p* < 0.01; *, *p* < 0.05. **(D)** MDA-MB-231 cells expressing mt-Keima were transduced the indicated sgRNAs and shRNAs. Cells were incubated with vehicle (DMSO) or BTZ (100nM) for 18hr prior to flow cytometry. Bar graphs represent mean +/- SEM from 3 independent experiments. Statistical analysis was performed using two-way ANOVA with Tukey’s post-test. ****, *p* < 0.0001; *, *p* < 0.05. **(E)** Presumptive model for endolysosomal regulation of mitophagy. Kinetic proofreading enforces the ultimate localization profile of BNIP3 despite limited targeting information, with the lysosome and proteasome serving as sinks that regulate available BNIP3. (Adapted from McKenna et al. 2020)

## DISCUSSION

Immense efforts have been directed toward understanding PINK1/Parkin-mediated mitophagy^2^. However, less is known about other mitophagy processes, including BNIP3-and NIX-mediated mitophagy. Early models for BNIP3-mediated mitophagy centered on its transcriptional control, particularly in response to hypoxic stress^4, 71^. Recently, these models have been appended to account for post-translational control by the ubiquitin-proteasome system^12–16, 56, 72^. Using an unbiased, genome-wide CRISPR screen, we similarly identified a role for the ubiquitin-proteasome system in regulating BNIP3, providing independent support for these models. However, prior reports do not fully account for BNIP3 dynamics in the cell.

As is documented for many autophagy receptors, lysosomal degradation of BNIP3 was presumed to be primarily through autophagy. Here, we demonstrate an alternative mode of BNIP3 degradation that is lysosome-mediated yet autophagy-independent. This alternative lysosomal delivery accounts for the vast majority of BNIP3’s lysosome-mediated turnover, even upon mitophagy induction. Consequently, lysosomal delivery of BNIP3 and/or NIX is an unexpectedly poor correlate for BNIP3/NIX-mediated mitophagy.

Our data indicate that the endolysosomal system functions independently of proteasomal regulation to further modulate levels and localization of BNIP3. When both mechanisms are disrupted, we see an additive increase in BNIP3 protein levels with a corresponding increase in mitophagy (Fig 4B, Fig 6B). Inhibition of ER insertion does not result in the overall accumulation of BNIP3 due to the compensatory effects of the proteasome. Regardless, the deletion of EMC components spatially restricts BNIP3 to the mitochondria, elevating mitophagy. In short, while BNIP3 can be cleared by parallel and partially compensatory quality control pathways, non-autophagic lysosomal degradation of BNIP3 is a strong post-translational modifier of BNIP3 function in both normoxic and hypoxic conditions.

With a new perspective on BNIP3 regulation, we took a structure-function approach to clarify the role of multiple conserved regions of BNIP3 including the LC3-interacting region (LIR), the BH3 domain and its adjacent ‘conserved region’, and the C-terminal TMD. The N-terminal LIR motif (ϕ-x-x-ψ) is a phospho-regulated motif governing the interaction of BNIP3 with ATG8-family proteins^6, 7^. As previously reported, we find that mutation of the LIR motif fully ablates BNIP3-mediated mitophagy. However, this region has no bearing on the lysosomal delivery of BNIP3, reinforcing BNIP3’s autophagy-independent flux. In contrast, BNIP3’s atypical BH3 domain has a modest effect on lysosomal delivery. Unlike canonical BH3 domains, this domain does not appear to function in cell death^61, 73–75^. Rather, it was recently implicated in the proteasome-mediated stability of BNIP3^16^. Our data support this interpretation, as truncation through the BH3 domain rendered BNIP3 resistant to proteasome inhibition by bortezomib (Fig. S4A). Continuous with the BH3 domain is an 11 amino acid conserved region of unknown function^76^. We find that truncation through this conserved region strongly disrupts the endolysosomal trafficking of BNIP3, leading to a mixed ER/mitochondria distribution (Fig. S4B). While we cannot exclude other functions for this region, we speculate that its conservation is a function of its role in the endolysosomal regulation of BNIP3. Finally, the C-terminal TMD of BNIP3 contains a GxxxG motif required for homodimerization^63, 77^. Disruption of this motif ablated the formation of SDS-resistant dimers but did not affect mitophagy, as measured by a highly quantitative mt-Keima assay. Similarly, overexpression of a chimera protein, BNIP3(Fis1^TMD^), induced mitophagy comparable to wild-type BNIP3, although BNIP3(Fis1^TMD^) was no longer subject to endolysosomal degradation. These data contrast with previous models, wherein oligomerization governs the activation of autophagy receptors. Formally, we cannot reject that 1) the soluble domain of BNIP3 provides sufficient self-association for mitophagy^78^ or 2) clustering of BNIP3 is driven through interaction with a soluble autophagy scaffold^79^. Yet, our data clearly indicate that the TMD of BNIP3 is dispensable for BNIP3-mediated mitophagy, contrary to early reports. Going forward, the ability to functionally separate the mitophagy and ER-trafficking activities of BNIP3 provides a foundation for future testing of more specific hypotheses regarding BNIP3 function *in vivo*.

More broadly, our findings have general implications for membrane protein quality control. Organelle identity and function are largely defined by the unique composition of each organelle’s constituent proteins. At first glance, the dynamic exchange of membrane proteins between organelles would appear paradoxical. However, growing evidence suggests that kinetically driven cycles of insertion and extraction–rather than a single, high-fidelity insertion event–best explain the observed, steady-state partitioning of many membrane proteins^28–33^. Perturbing this cycle results in the aberrant accumulation of TA proteins at incorrect membranes. Extending these observations, we find constitutive delivery of BNIP3, a model TA protein, to the ER in the absence of any genetic perturbation. In this context, BNIP3 delivery is strongly dependent on the EMC, with the GET complex playing a lesser role. This is congruent with the observation that mitochondrial TA proteins and EMC substrates possess similarly hydrophilic TMDs, although previous studies suggest the GET pathway can intercede when confronted with a significant buildup of non-optimal TA substates^30, 80^.

Mitochondrial TA proteins that mislocalize to the ER are recognized by ATP13A1 (Spf1 in yeast), an ER-resident ATPase functionally analogous to ATAD1/Msp1 in the OMM^29, 33^. Supporting its role as a TA protein extractase, deletion of ATP13A1/Spf1 results in the accumulation of mitochondrial TA proteins on the ER^81–84^. Why, then, might cells require an alternative ER clearance system à la the endolysosomal pathway employed for BNIP3? An emerging paradigm for TA protein extractases is that orphan TA proteins are preferred substrates^84, 85^. This includes excess or mislocalized TA proteins that fail to incorporate into stable, higher-ordered complexes. Because BNIP3 self-associates, we anticipate that the formation of a stable homodimer renders BNIP3 resistant to ATP13A1-mediated extraction and necessitates an alternative quality control mechanism. At the same time, BNIP3 dimerization is strictly required for lysosomal delivery. Therefore, we propose that self-association enforces a switch between proteasomal and lysosomal degradation routes. In further support of this model, we found accelerated clearance of BNIP3 monomers at both mitochondria and the ER, in a strictly proteasome-dependent manner (Fig 5I). The role of ATAD1 and ATP13A1 in destabilizing and/or shuttling these BNIP3 monomers is beyond the scope of this work but will be an important area of future study. In total, our results support a model where extraneous or mislocalized TA monomers are degraded by the UPS, while dimerization leads to stable protein complexes that are cleared from the ER through trafficking to lysosomes. In such a model, BNIP3 dimers present in the OMM are resistant to both forms of quality control, thus explaining the observed steady-state localization of BNIP3 in the OMM. While other groups have speculated such a model,^29, 33^ we provide direct evidence using an endogenous TA protein, BNIP3. Thus, we directly implicate endosomal trafficking and lysosomal degradation in the canon of quality control pathways that support the proper localization of TA membrane proteins.

BNIP3 has been implicated in a variety of physiological processes not considered here^52, 86–93^. Consequently, the full physiological implications of BNIP3 regulation will be an important area of continued study. For instance, a tumor-suppressor function for BNIP3 has been suggested that is independent of its role as a BH3-containing protein^88^. Correspondingly, transcriptional repression of BNIP3 is associated with several cancer types^94^. Given the extent to which post-translational mechanisms impinge upon BNIP3 function, we anticipate that post-translational control of BNIP3 may similarly be exploited by cancerous cells to restrict BNIP3. Since transcriptome-level analyses are blind to this level of regulation, BNIP3’s role in tumor progression is likely underestimated.

We note that BNIP3-mediated mitophagy is commonly associated with stress conditions, particularly where hypoxia is a factor as in ischemia/reperfusion injury^86, 95, 96^. In contrast, NIX generally is implicated in mitophagy during cellular differentiation programs^97–103^. Future efforts will be needed to further delineate the differential utilization of these highly related mitophagy receptors. However, this utilization trend is generally consistent with the relative responsiveness of BNIP3 and NIX to proteostatic collapse. Going forward, it will also be important to fully consider the implications of proteostatic collapse on mitophagy. For example, bortezomib-induced peripheral neuropathy (BIPN) is a common and painful side effect of bortezomib use as a chemotherapeutic agent^104, 105^. While the underlying mechanism of BIPN remains a matter of debate, the mitochondrial dysregulation associated with BIPN makes BNIP3-induced mitophagy an intriguing therapeutic candidate.

## MATERIALS AND METHODS

### Antibodies

For immunoblotting (IB), all primary antibodies are used at a 1:1,000 dilution, unless stated otherwise. Secondary antibodies are used at a 1:10,000 dilution. For immunofluorescence (IF): primary antibodies were diluted 1:100 and secondary antibodies were used 1:1000. The follow primary antibodies were used: mouse anti-SQSTM1/p62 (ab56416, Abcam), rabbit anti-NDP52 (9036, CST), rabbit anti-ATG9A (13509S, CST), mouse anti-GFP (118114460001, Sigma), rabbit anti-BNIP3 (44060S, CST), rabbit anti-BNIP3L/NIX (12396S, CST), anti-V5 Tag (13202, CST), rat anti-tubulin (sc-53030, Santa Cruz Biotechnology), mouse anti-tubulin (3873S, CST), and mouse anti-EMC3/TMEM111 (67205-1-lg, Proteintech). The following secondary antibodies were used for (IB): goat anti-mouse IgG(H+L) IRDye 680LT (926-68020, LI-COR), goat anti-rabbit IgG(H+L) IRDye800CW (926-32211, LI-COR); secondary antibodies (IF): goat anti-rabbit IgG(H+L) Alexa Fluor Plus 647 (A32733, Invitrogen), goat anti-mouse IgG(H+L) Alexa Fluor Plus 647 (A32728, Invitrogen).

### Vectors

The Brunello knockout pooled library was a gift from David Rootand John Doench (Addgene #73178). psPAX2 was a gift from Didier Trono (Addgene plasmid # 12260). pCMV-VSV-G was a gift from Bob Weinberg (Addgene plasmid #8454). lentiCRISPRv2puro was a gift from Brett Stringer (Addgene plasmid #98290). lentiGuide-puro was a gift from Feng Zhang (Addgene plasmid #52963). pFUGW-EFSp-Cas9-P2A-Zeo (pAWp30) was a gift from Timothy Lu (Addgene plasmid #73857). pLenti CMV GFP Puro (658-5) was a gift from Eric Campeau & Paul Kaufman (Addgene plasmid #17448). mito-BFP was a gift from Gia Voeltz (Addgene # 49151). pGW1-mCherry-EGFP-PIM was a gift from Lukas Kapitein (Addgene plasmid #111759). pHAGE-mt-mKeima was a gift from Richard Youle (Addgene plasmid #131626). pLKO.1 hygro was a gift from Bob Weinberg (Addgene plasmid # 24150) Other vectors generated during this study are available upon request.

### Chemicals and Reagents

The following chemicals and reagents were used in this study: 2-Deoxy-D-glucose (D8375-1G, Sigma), 2-mercaptoethanol (BME) (M6250-100ML, Sigma), agar (A10752, AlphaAesar), agarose (16500500, Thermo Fisher), ampicillin (A9518-25G, Sigma), Bafilomycin A1 (11038, Caymen Chemical), Beta-glycerophosphate (35675-50GM, Sigma), blasticidin (ant-bl-1, Invivogen), Bortezomib (10008822, Caymen Chemical), Brefeldin-A1 (11861, Caymen Chemical), CB-5083 (19311, Caymen Chemical), dimethyl sulfoxide (C833V25, Thomas Scientific), EDTA (EDS-500G, Sigma), glycerol (G2025-1L, Sigma), HEPES (H3375-1KG, Sigma), hygromycin (ant-hg-1, Invivogen), kanamycin (BP906-5, FisherSci), MLN-4924 (15217, Caymen Chemical), MLN-7243 (30108, Caymen Chemical), plasmocin (ant-mpp, Invivogen), Phusion High-Fidelity DNA poly-merase (M0530L, NEB), PIK-III (17002, Caymen Chemical), polybrene (H9268-5G, Sigma), potassium chloride (P217-500, FisherSci), puromycin (ant-pr-1, Invivogen), sodium chloride (6438, FisherSci), sodium deoxycholate (97062-028, VWR), sodium dodecyl sulfate (SDS)(74255-250G, Sigma), sodium fluoride (S6776-100G, Sigma), sodium orthovanadate (450243-10G, Sigma), sodium pyrophosphate decahydrate (221368-100G, Sigma), Taq DNA ligase (M0208L, NEB), TERGITOL Type NP-40 (NP40S-100ML, Sigma), Tris base (T1378-5KG, Sigma), TritonX-100 (T9284-500ML, Sigma), tryptone (DF0123-17-3, FisherSci), Tween-20 (BP337-500, FisherSci), T5 exonuclease (M0363S, NEB), yeast extract (BP1422-2, FisherSci), and zeocin (ant-zn-1, Invivogen).

### Tissue Culture

All mammalian cells were grown in a standard water-jacketed incubator with 5% CO_2._ MDA-MB-231, U2OS, HEK293T, MDA-MB-453 all grown in DMEM media (45000-304, Corning) with 10% FBS (26140079, Gibco) and 1X penicillin/streptomycin (15140122, Thermo Scientific). K562 cells were grown in IMDM media (45000-366, Corning) with 10% FBS and 1X penicillin/streptomycin. All mammalian cells were acquired from American Type Culture Collection (ATCC). Plasmocin prophylaxis (1:500) was used when generating of new stable cell line. All cells were maintained below an 85% confluency and passaged less than 25 times. For passaging, cells are trypsinized with 0.25% Trypsin-EDTA (25200114, Thermo Scientific). For hypoxia incubation, cells were incubated in a humidified Baker Ruskinn InvivO_2_ (I400) hypoxia chamber at 1% O_2_ and 5% CO_2_ for the indicated times. Puromycin (2*µ*g/mL), blasticidin (5*µ*g/mL), and zeocin (50*µ*g/mL) were added when necessary for selection. 1X Hanks’ Balanced Salt Solution (HBSS) (45000-456, Corning) was used to wash cells when passaged.

### Generation of gene knockout cell line

Sequences for sgRNAs targeting genes of interest were extracted from the Brunello library and cloned into the indicated vectors as outlined above under “sgRNA oligonucleotide ligation protocol”. HEK293T and MDA-MB-231 cells were transfected with the resulting vectors. Single cell sorting was used to isolate individual clones. Knockouts of expanded clones was confirmed by immunoblotting.

### Molecular cloning and bacterial transformation

PCR inserts were amplified using Phusion High-Fidelity DNA polymerase (M0530L, NEB). Amplification primers were designed with a 30 base pair overlap with the linear ends of restriction-digested recipient vectors. Linearized vector backbones were dephosphorylated by calf intestinal phosphatase (M0290S, NEB). All inserts and vectors were purified from a 0.9% agarose gel prior to isothermal assembly (D4002, Zymo Research). 50ng of linearized vector DNA was combined with isomolar amounts of purified insert(s). 2.5*µ*L DNA mix was incubated with 7.5ll isothermal assembly master mix at 50°C for 20 min. Product of the isothermal assembly reaction was transformed into NEB Stable cells (C3040H, NEB). Transformed cells were plated on plates of LB media (10 g/L tryptone, 5 g/L yeast extract, 5 g/L NaCl) containing 1.5% agar, 100μg/mL ampicillin or 50μg/mL kanamycin were included in bacterial cultures, where appropriate. All cultures and plates were grown overnight at 34°C. Overnight cultures were pelleted at 3,000g for 10 min and plasmid DNA was purified using a Qiagen miniprep kit (27106, Qiagen). Sequences were verified by Sanger sequencing (Eton Bioscience Inc).

### sgRNA oligonucleotide ligation

Oligonucleotides were ordered from Thermo Fisher. For sgRNA cloning, oligos were ordered in the following format: Forward: 5’-CACCGNNNNNNNNNNNNNNNNNNNN-3’; Reverse: 5’-AAACNNNNNNNNNNNNNNNNNNNNC-3’. For shRNA cloning, oligos were ordered in the following format: Forward: 5’-CCGGNx48TTTTTG-3’; Reverse: 5’-AATTCAAAAANx48-3’. 50pmol of each oligo were mixed in a 25*µ*L reaction and phosphorylated with T4 polynucleotide kinase (M0201S, NEB). Reactions were performed for 30 min at 37°C in 1X T4 DNA ligase buffer (B0202S, NEB). Phosphorylated oligos were annealed by heating for 5 min at 95°C and slow cooling (0.1°C/s). 2μl of diluted (1:100) oligo mix was ligated into 20 ng BsmBI-digested vector (pLentiGuide-puro or pLenti-CRIPSR v2), or AgeI/EcoRI-digested vector (pLKO.1 hygro for shRNA), using T4 DNA ligase (M0202S, NEB). Ligation reaction was done at room temperature for 15 min prior to bacterial transformation. All small hairpin and sgRNA sequences are listed in Table S2.

### Flow Cytometry

Cells were trypsinzed, centrifuged, and resuspended in cold 1X HBSS and filtered through a 41-*µ*m nylon mesh prior to flow analysis. All flow cytometry data was collected on a Beckman Coulter CytoFLEX flow cytometer. Data was analyzed using FlowJo v10 and R Studio. At least 10,000 cells were collected for all samples.

### Lentiviral generation

Lentivirus was generated using HEK293T cells with Lipofectamine 3000. Cells were seeded in Opti-MEM media, containing 5% FBS and no antibiotics, overnight for 80% confluency. Cells were transfected with packaging vectors pVSV-G and pSPAX2, along with expression construct at a 1:4:3 ratio, scaled accordingly. Opti-MEM media was refreshed 6hr after transfection. Supernatant containing virus was collected at 24-and 48-hr post-transfection and pooled together. Virus was cleared by centrifugation for 15 min at 1000 g and aliquoted to avoid freeze-thaw cycles.

### Viral transduction

Cells were incubated in respective media containing 8*µ*g/mL polybrene (1:1000 dilution) with virus. If adherent, cells were tryspinized and allowed to re-adhere with media containing polybrene and virus. Transduction were left overnight, and virus-containing media was exchanged in the morning with fresh media lacking polybrene. Transduced cells were allowed to recover in fresh media for 24hr prior to antibiotic selection.

### Transient transfection

Cells were seeded at approximately 75% confluency in Opti-MEM reduced media supplemented with 5% FBS no antibiotics and allowed to adhere overnight. Cells were transfected using Lipofectamine 3000 reagent (L3000008, Life Technologies), according to the manufacturer’s protocol. Cells were left in Lipofectamine reaction for 6hr before a fresh Opti-MEM media exchange. Cells were analyzed 24hr post-transfection.

### Gel electrophoresis and immunoblotting

Cells are resuspended and washed once in 1X HBSS prior to lysis. Cells are lysed for 20 min on ice in lysis buffer (50mM HEPES pH 7.4, 40mM NaCl, 2mM EDTA, 1% Triton X-100, 2X complete protease inhibitor tablet (5056489001, Sigma)). Lysates were cleared by centrifugation for 8 min at 1000 g, using supernatants as sample input. Total protein level was normalized using a BCA protein assay (23225, Thermo Scientific) and adjusted with lysis buffer. Normalized lysate samples were boiled at 65 degrees for 10 min in 1X (final concentration) Laemmli Loading Buffer (3X stock: 189mM Tris pH 6.8, 30% glycerol, 6% SDS, 10% beta-mercaptoethanol, bromophenol blue). Gel electrophoresis was performed at 175V for 60 min in Novex 4-20% Tris-Glycine gels. Protein gels were transferred for 60 min to 0.2*µ*m PVDF membranes (ISEQ00010, Sigma) using the semi-dry Trans-Blot Turbo Transfer system (Bio-Rad). Membranes were blocked for at least 30 min in 5% Milk in 1X TBST (MP290288705, Fisher Scientific). Primary antibodies were diluted in 5% Milk in TBST and incubated on membrane overnight at 4°C. After overnight primary incubation, membranes were washed three times in 1X TBST for 5 min. Secondary antibodies were diluted in Intercept^TM^ (TBS) Blocking Buffer (927-60003, LI-COR) and incubated on membrane for 1hr at room temperature. After secondary incubation, membranes were washed twice in 1X TBST and last wash was done in 1X TBS (no Tween). All membranes were imaged on LI-COR Odyssey CLx dual-color imager and band intensities were quantified on LI-COR analysis software Image Studio Lite.

### Mito-Keima assays

For BFP-BNIP3 overexpression, MDA-MB-231 cells stably expressing mt-Keima reporter were transduced following normal viral transduction. Transduced cells were either treated with vehicle (DMSO) or Baf-A1 (100nM) after 24hr post-transduction for 18hr and analyzed by flow cytometry 48 hr post-transduction. For drug treatment, MDA-MB-231 cells stably expressing mt-Keima reporter were treated with respective drug for 18 to 24-hr prior to flow cytometry. Baf-treated and non-transduced samples served as gating controls for all mt-Keima flow analysis.

### Measuring oxygen consumption

Oxygen consumption and glycolytic rates were analyzed using the Seahorse XF96 system. Cells were seeded on Seahorse XF96 cell culture microplates (101085-004, Agilent) at a density of 0.25 x 10^5^ density per well in DMEM media supplemented with 10% FBS and no antibiotic selection 24hr prior to analysis. DMEM media was exchanged with Mito Stress XF DMEM media and incubated for 35min prior to test. The Mito Stress Test (103015-100, Agilent) was performed the following day, using the manufacturer’s protocol (Injection 1: Oligomycin 1.5*µ*M; Injection 2: FCCP 1*µ*M; Injection 3: Rotenone/Antimycin A 0.5*µ*M; Injection 4: 2-Deoxy-D-glucose 50mM). Respiration and glycolytic rates were calculated based on manufacturer’s protocol. The Seahorse XF96 analyzer from the Immune Monitoring and Flow Cytometry Shared Resource (DartLab) was used. All Seahorse data was normalized by cellular lysis using RIPA lysis buffer (150mM NaCl, 50mM Tris-HCl pH 8, 0.5% sodium deoxycholate, 0.1% SDS, 1% TERGITOL Type NP-40 solution, 2X complete protease inhibitor tablet) in the microplate and performing a BCA protein assay.

### Artificial Dimerization Assay

The DmrB inducible dimer domain was subcloned from pGW1-mCherry-EGFP-PIM (addgene #111759) to the N-terminal cytosolic portion of BNIP3. DmrB constructs were packaged in lentivirus and transduced into cells. Transduced cells were exchanged with fresh media and allowed to recover for 24hr. On day 2 post-transduction, cells were exchanged with fresh media containing B/B homodimerizer (500nM) (Takara Bio, #635059) and/or Baf-A1 (100nM) and/or vehicle (DMSO) control for 6hr prior to flow cytometry analysis.

### *In vitro* dephosphorylation assay

Cells were transduced with lentivirus for expression of V5-BNIP3 variants. Cells were lysed in high salt/IP lysis buffer (50mM HEPES pH 7.4, 150mM NaCl, 2mM EDTA, 1% Triton X-100, 2X complete protease inhibitor tablet (5056489001, Sigma)). Lysates were cleared by centrifugation. 25*µ*L of Magnetic V5-Trap bead slurry (v5tma-10, Chromotek), per lysate sample, was washed once with IP lysis buffer and incubated with cleared lysates for 40min at 4°C on end-over-end rotator. Pull-down flow through was collected after bead-lysate incubation. Bead slurry was divided in 3 and washed five times with IP lysis buffer. All three bead samples were moved to PMP buffer (P0753S, NEB), corresponding to PMP buffer only, PMP with Lambda Protein Phosphatase, and PMP with Lambda Protein Phosphatase (P0753S, NEB) with Phosphatase inhibitor cocktail (4X: 5mM NaF, 1mM orthovanadate, 1mM pyrophosphate, 1mM glycerophosphate). For 50*µ*L reactions, the following volumes were used: 5*µ*L of 10X PMP buffer, 5*µ*L of MnCl_2_ (10mM), 0.75*µ*L of Lambda Protein Phosphatase, and 4X Phosphatase inhibitor cocktail. Dephosphorylation reaction was done for 30min at 30°C prior to sample denaturing.

### Immunofluorescence and live cell microscopy

Cells were seeded on glass bottom dishes (07-000-235 and NC0832919, Fisher Scientific) at approximately 70% confluency and allowed to adhere overnight. Cells were washed twice in warm 1X HBSS and fixed for 15min in 4% paraformaldehyde (PFA) made from fresh 16% PFA (#15710, Electron Microscopy Sciences) diluted with 1X HBSS. Cells were blocked and permeabilized at room temperature for 1h in Intercept^TM^ (TBS) Blocking Buffer (927-60003, LI-COR) plus 0.3% Triton X-100, then washed once in 1×HBSS. Primary anti-body was diluted in (1:100), and cells were incubated in Intercept^TM^ (TBS) Blocking Buffer of primary antibody solution overnight at 4°C. After incubation, cells were washed 3×5 min in 1×HBSS. Secondary antibody was diluted to 1:1,000 in Intercept^TM^ (TBS) Blocking Buffer, and cells were incubated in secondary antibody solution for 45 min at room temperature. After incubation, cells were washed 3×10min in 1×HBSS, stained with a 1:10,000 dilution of Hoechst 33342 (H3570, Thermo Fisher) for 5min, and washed once more in 1×HBSS before storage or imaging. For living cell imaging, cells were seeded on glass bottom dishes at approximately 70% confluency and imaged in DMEM with no phenol red (21-063-029, Fisher Scientific) supplemented with 10% FBS.

### Confocal microscopy

All fluorescent images were obtained using a Nikon Eclipse Ti-E inverted microscope stand that has a Yokogawa, two-camera, CSU-W1 spinning disk system with a Nikon LU-N4 laser launch. All images were obtained on 100X PlanAPO objective lens.

### Image analysis

Image intensities were modified linearly and evenly across samples per experiment. All images represent a single plane on acquired Z-stack and processed in ImageJ. For co-localization analysis, the Coloc2 plugin on ImageJ was used. Image intensities were modified and threshold for the two channels of interest. Cellular segmentation in images were done manually to obtain region of interests (ROIs). ROIs underwent Coloc2 analysis.

### Library propagation

Brunello library was purchased from Addgene (#73178). Library (50ng) was electroporated into 25*µ*L Endura electrocompetent cells (60242-2, Lucigen). Cells from eight electroporations were pooled and rescued in 8mL of rescue media for 1hr at 37. 8mL of SOC media (2% tryptone, 0.5% yeast extract, 10mM NaCl, 2.5mM KCl, 10mM MgCl_2_, 10mM MgSO_4_, and 20mM glucose) was added to electroporated cells and 200*µ*L of the final volume was spread onto 10cm LB plates containing 50*µ*g/mL carbenicillin (80 LB plates total). Cells were manually scraped off plates to perform a plasmid DNA purification using GenElute Megaprep kit (NA0600-1KT, Sigma).

### Library lentiviral generation

Lentivirus for Brunello library was generated by lipofection of HEK293T cells with 5*µ*g psPAX2, 1.33*µ*g pCMV-VSV-G, and 4*µ*g of library vector per 10 cm plate of HEK293T at 85% confluency. Low-passage HEK293T cells were grown in OptiMEM containing 5% FBS and no antibiotics. OptiMEM media was exchanged 6hr after transfection. Supernatants containing virus were collected at 24 hr post-transfection, replenished, and collected again at 48hr. Supernatants were pooled and cleared by centrifugation for 15 mins at 1000 g. Viral titer was quantified using the Lenti-X™ qRT-PCR Titration Kit (Takara Bio, #631235), according to manufacturer’s protocol.

### Transduction and cell growth

For CRISPR screening experiments, MDA-MB-231 cells were passaged to maintain cell density between 80-90% confluency in 10cm dishes. Cells were propagated in DMEM+10% FBS + pen/strep + appropriate antibiotics (Blasticidin 5μg/ml, zeocin 50μg/ml) until 100 million cells were obtained (approximately 8–10 days). 100 million cells were trypsinized and resuspended in DMEM + 10% FBS + 8μg/mL polybrene. An MOI of 0.4 was used to minimize multiple infection events per cell. Date of infection was day 0. Cells were infected over-night and exchanged into fresh media. After 24 h, 2μg/ml puromycin was added. Cells were expanded to 15cm dishes in puromycin. Cells were removed from puromycin 1 day prior to sorting and at day 8, 11, 12, cells were sorted for red:green fluorescence, sorting 200 million cells each day. 50 million unsorted cells were collected and processed as input. The top and bottom 30% of cells (based on Red:Green ratio) were taken. Cell sorting was performed using a Sony SH800 cell sorter. Cells were pelleted and stored at 80°C until processing.

### CRISPR Screen Processing

Genomic DNA was purified from collected cells using the NucleoSpin Blood XL kit (740950.1, Macherey Nagel) according to the manufacturer’s instructions. sgRNA sequences were amplified from total genomic DNA using a common pool of eight staggered-length forward primers. Unique 6-mer barcodes within each reverse primer allowed multiplexing of samples. Each 50μL PCR reaction contained 0.4*µ*M of each forward and reverse primer mix (Integrated DNA Technologies), 1×Phusion HF Reaction Buffer (NEB),0.2 mM dNTPs (NEB), 40 U/ml Phusion HF DNA Polymerase (NEB), up to 5lg of genomic DNA, and 3% v/v DMSO. The following PCR cycling conditions were used: 1×98°C for 30 s; 25× (98°Cfor 30 s, 56°C for 30 s, 63°C for 30 s); 1×63°C for 10 min. The resulting products were pooled to obtain the sgRNA libraries. The pooled PCR products were size selected between 0.60×and 0.85×magnetic bead slurry as outlined by DeAngelis al(1995). Library purity and size distribution was measured on a Fragment Analyzer instrument (Agilent) and quantified fluorometrically by Qubit. Libraries were pooled in equimolar ratios and loaded at 2.5 pM on to a NextSeq500 High Output 75cycle run. 2% PhiX spike in was included as an internal control for sequencing run performance. Data were demultiplexed into fastq files using Illumina bcl2fastq2v2.20.0.422.

### NGS data analysis

The 5’ end of Illumina sequencing reads was trimmed to 50-CACCG-30using Cutadapt (Martin, 2011). The count function of MAGeCK (version 0.5.9) was used to extract read counts for each sgRNA (Li et al, 2014). Trimmed fastq files are available on Mendeley Data using the DOIs listed in key resources table. The RRA function was used to compare read counts from cells displaying increased and decreased Red:Green ratios (Li et al, 2015). The output included fold change, rank, and p-value. Average fold change scores (across 3 experimental replicates), and p-values can be found in Table S1.

### Statistical analysis

All statistical analysis was performed using Prism 8 (GraphPad). All statistical tests are indicated in the relevant figure legends. All tests were two-tailed with *P < 0.05* as the threshold for statistical significance. Number of replicates (n) used for each experiment and statistical test are indicated in the relevant figure legend.

## ACKNOWLEDGEMENTS

We would like to thank former lab member Amelia Ohnstad for technical assistance. We would like to thank Dr. Michael Ragusa and lab members for providing insightful feedback. We would like to thank Dr. Robert Cramer and Kaesi Morelli for assistance with the hypoxia chamber. We would like to thank Vladimir Denic, Michael Ragusa, and Charles Barlowe for critical reading of the manuscript. This work is supported by the National Institutes of Health General Medical Sciences (R35GM142644 to CJS, F31GM143849 to JMD). We would like to thank Ann Lavanway for light microscopy expertise. We would like to thank the Institute for Biomolecular Targeting (bioMT) core at Dartmouth supported by P20GM113132. We would like to thank the Genomics Shared Resource and the Immune Monitoring and Flow Cytometry Shared Resource (DartLab) supported by P30CA023108 to the Dartmouth Cancer Center.

## SUPPLEMENTAL FIGURE LEGENDS

**Figure S1, related to figure 1.**
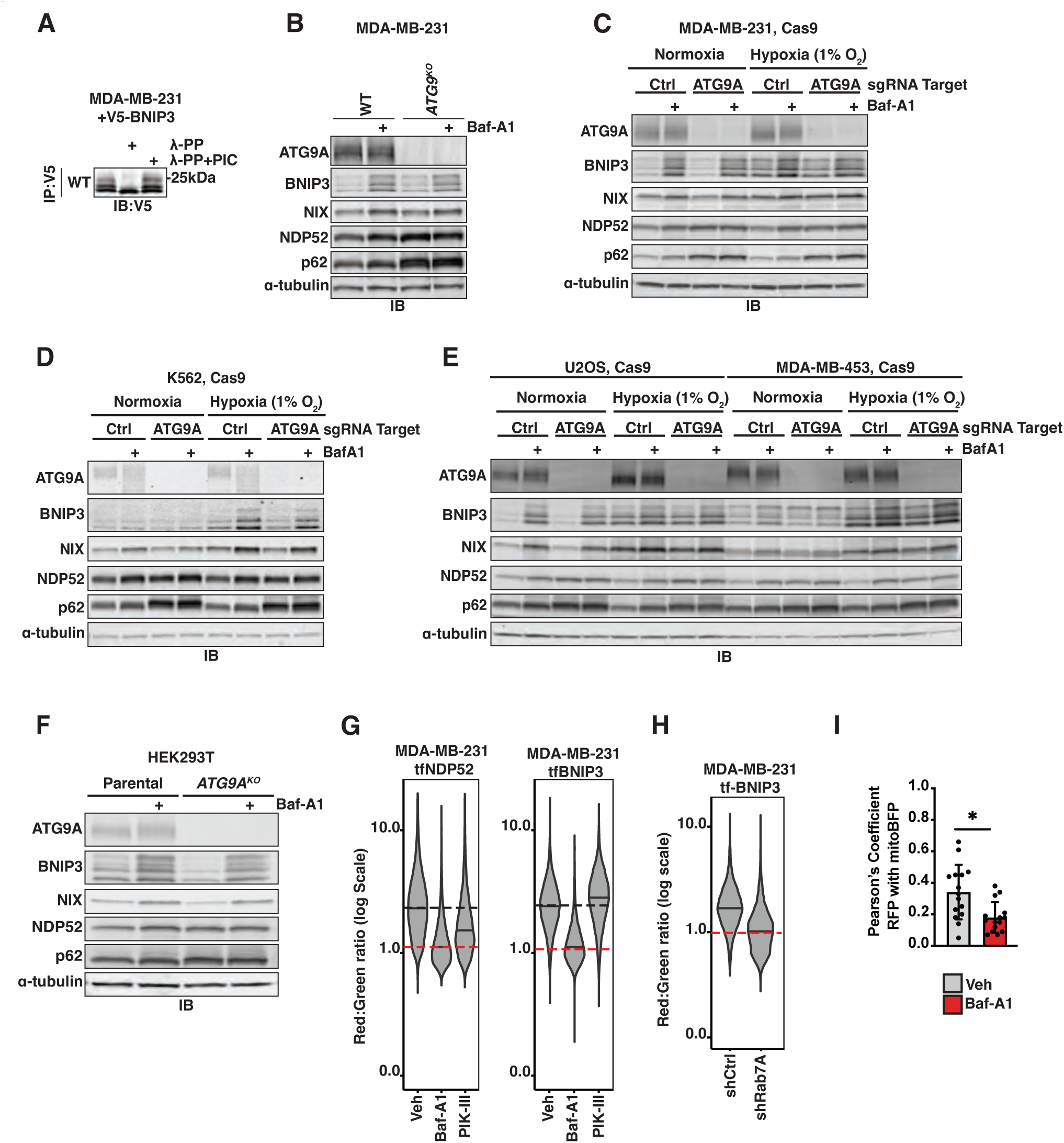
**(A)** MDA-MB-231 cells were transduced with V5-BNIP3 variants and lysed 48hr post-transduction. The V5 epitope was immunoprecipitated from extracts and treated with buffer alone (lane 1), lambda phosphatase (PP, lane 2), or lambda phosphatase with phosphatase inhibitor cocktail (PIC, lane 3). **(B)** Immunoblotting of MDA-MB-231-derived extracts from wild-type (WT) and *ATG9^KO^*clonal knockout cells. Where indicated, cells were treated with Baf-A1 (100nM) for 18hr. **(C-E)** Immunoblotting of MDA-MB-231, K562, U2OS, MDA-MB-453-derived extracts from cells expressing Cas9 and the indicated sgRNA. Cells were subjected to normoxia or hypoxia and/or Baf-A1 treatment (100nM) for 18hr where indicated. **(F)** Immunoblotting of extracts derived from parental HEK293T and clonal *ATG9^KO^* knockout cells. Where indicated, cells were treated with Baf-A1 (100nM) for 18hr. **(G)** Violin plots of MDA-MB-231 cells expressing either the tf-NDP52 or tf-BNIP3 reporter. Cells were treated with DMSO or Baf-A1 (100nM) or PIK-III (10*µ*M) for 18h before being analyzed by flow cytometry for red:green ratio. Median values for each sample are identified by a black line within each violin. The red dotted line across all samples corresponds to red:green ratio of maximally inhibited conditions (Baf-A1) (n > 10,000 cells). **(H)** Violin plots of MDA-MB-231 cells expressing tf-BNIP3 transduced with either a control small hairpin RNA (shCtrl) or an shRNA targeting Rab7 (shRab7). Cells were analyzed for red:green ratio 8 days post-transduction. Red dotted line (=1) corresponds to the theoretical maximum inhibition of red:green ratio **(I)** Quantification of Pearson’s correlation coefficients from cells in Fig 1D. Correlation of RFP to mitoBFP (reflective of mitochondrial localization) was calculated using Coloc2. Bar graphs represent mean +/- SEM. Each data point represents a single cell. n = 15 cells. Statistical analysis was performed using an unpaired Student’s t-test. *, *p* < 0.05.

**Figure S2, related to figure 3.**
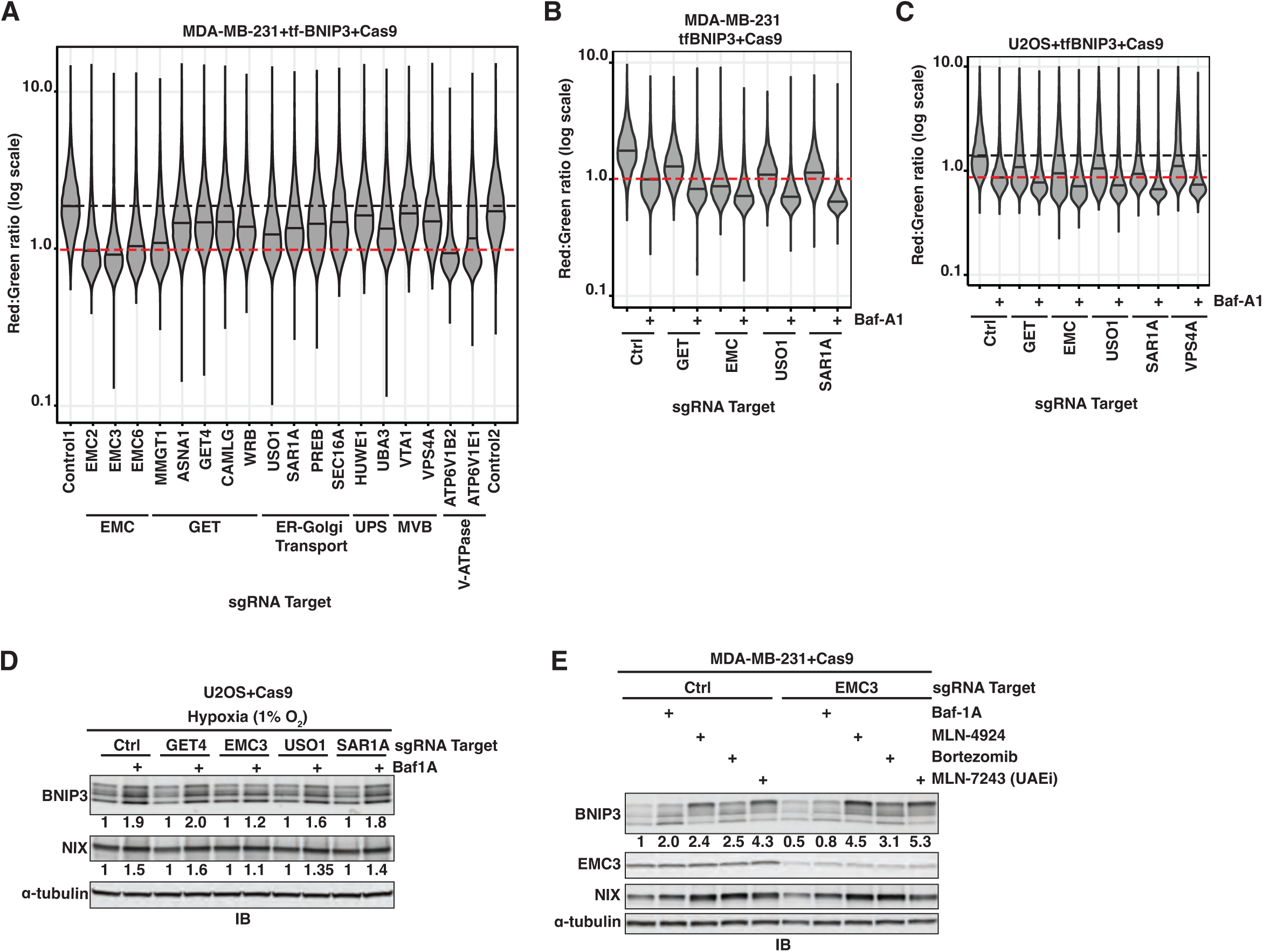
**(A)** MDA-MB-231 cells expressing tf-BNIP3 and Cas9 were transduced with the indicated sgRNAs. Red:green ratio was analyzed by flow cytometry on day 8 post-transduction. Median values for a non-targeting control (sgControl1) are identified by a dashed black line. The red dotted line across all samples corresponds to a red:green ratio of 1, the theoretical minimum (n > 10,000 cells). **(B)** MDA-MB-231 cells expressing tf-BNIP3 and Cas9 were transduced with the indicated sgRNAs. Red:green ratio was analyzed by flow cytometry on day 8 post-transduction. The red dotted line across all samples corresponds to red:green ratio of Baf-A1-treated control (Ctrl) cells (n > 10,000 cells). **(C)** U2OS cells expressing tf-BNIP3 and Cas9 were transduced with the indicated sgRNAs. Red:green ratio was analyzed by flow cytometry on day 8 post-transduction. The red dotted line across all samples corresponds to red:green ratio of Baf-A1-treated control (Ctrl) cells (n > 10,000 cells). **(D)** Immunoblotting of U2OS-derived extracts expressing Cas9 that were transduced with the indicated sgRNAs. On day 8 post-transduction, cells were treated with Baf-A1 (100nM) and subjected to hypoxia for 18hr prior to lysis. **(E)** Immunoblotting of MDA-MB-231-derived extracts expressing Cas9 that were transduced with the indicated sgRNAs. Cells were treated with Baf-A1 (100nM), MLN-4924 (1*µ*M), Bortezomib (100nM), or MLN-7243 (an inhibitor of the ubiquitin activating enzyme [UAE], 1*µ*M) for 18hr on day 8 post-transduction, prior to lysis.

**Figure S3, related to figure 4.**
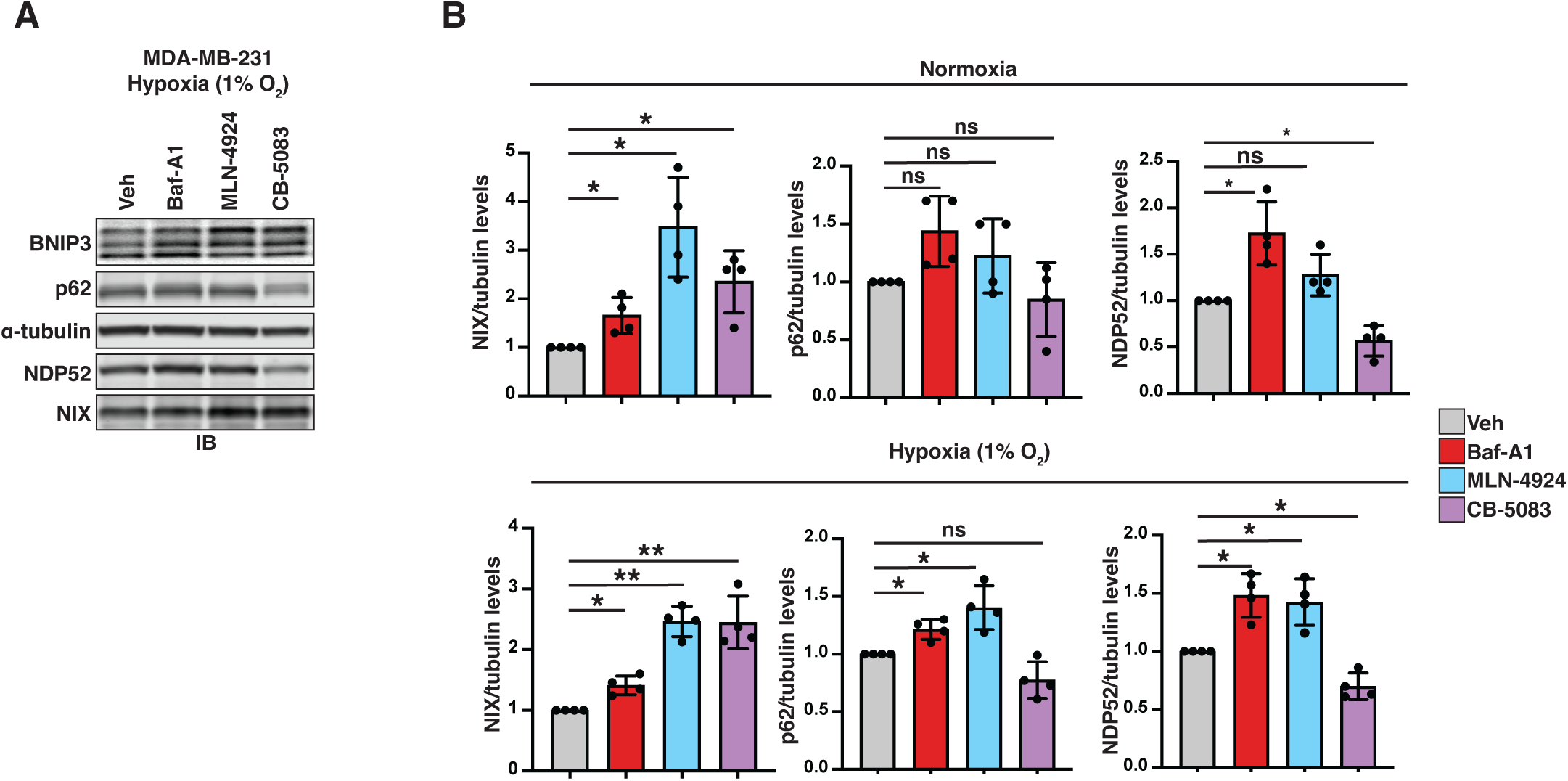
**(A)** Representative image of one biological replicate quantified in Fig 4A. Immunoblotting (IB) of MDA-MB-231-derived extracts from cells treated with vehicle (DMSO), Baf-A1 (100nM), MLN-4924 (1*µ*M), CB-5083 (1*µ*M) and subjected to hypoxia for 18h. **(B)** Quantification of protein accumulation from Fig 4D and Fig S3A. Bar graphs represent mean +/- SEM from 4 independent experiments. All protein levels were normalized to α-tubulin. Statistical analysis was performed using a one-sample t-test to the normalized control. **; *p* < 0.01; *; *p* < 0.05; *ns*, not significant.

**Figure S4, related to figure 5.**
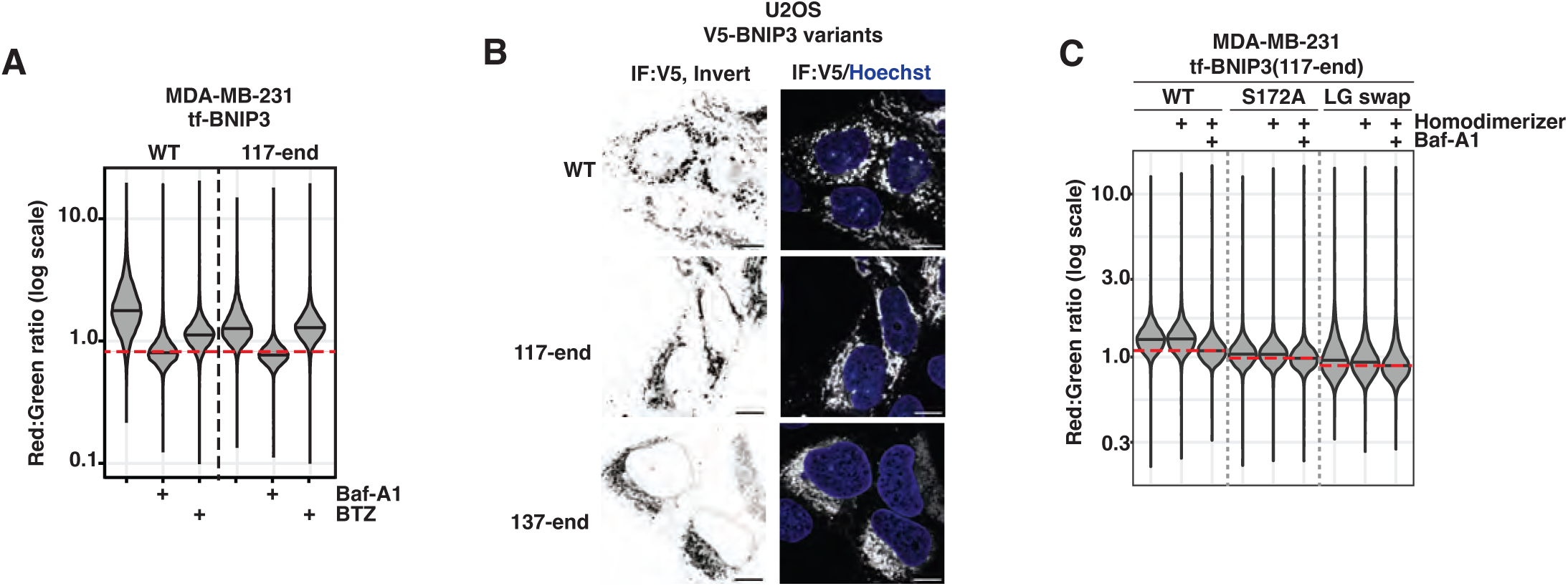
**(A)** MDA-MB-231 cells were transduced with the indicated tf-BNIP3 variants. Red:green ratio was analyzed by flow cytometry 48hr post-transduction. Cells were treated with vehicle (DMSO), Baf-A1 (100nM), or BTZ (100nM) for 18hr prior to performing flow cytometry. The red dotted line across each sample group corresponds to the maximum inhibition red:green ratio of the wild-type (WT) Baf-A1-treated sample (n > 10,000 cells). **(B)** Representative confocal micrographs of U2OS cells transduced with V5-BNIP3 variants. 48hr post-transduction, cells were fixed and immunostained for the V5 epitope. Hoechst stain was used for nuclear staining. Scale bar is 10*µ*m. **(C)** MDA-MB-231 cells were transduced with the indicated tf-BNIP3^117^–^end^ variants. Red:green ratio was analyzed by flow cytometry 48h post-transduction. Cells were treated with Baf-A1 (100nM) and/or B/B homodimerizer (0.5 *µ*M) for 6h prior to performing flow cytometry. Median values for each sample are identified by a black line within each violin. The red dotted line across each sample corresponds to cells inhibited with Baf-A1 (n > 10,000 cells).

**Figure S5, related to figure 6.**
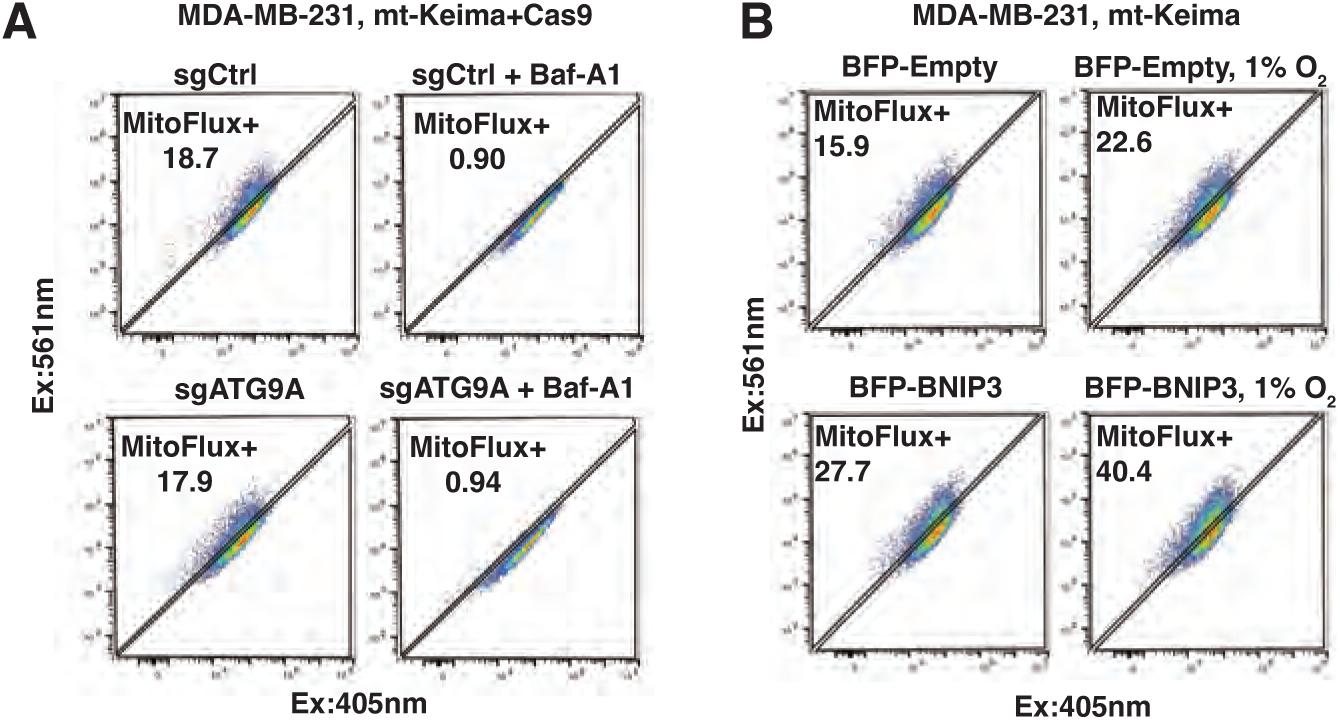
**(A)** MDA-MB-231 cells expressing mt-Keima were transduced with either a non-targeting sgRNA (sgCtrl) or sgATG9A. On day 8 post-transduction, cells were incubated with vehicle (DMSO) or Baf-A1 (100nM) for 18hr and assessed by flow cytometry. **(B)** MDA-MB-231 cells expressing mt-Keima were transduced with BFP-BNIP3. At 24hr post-transduction, cells were incubated in normoxic or hypoxic conditions for 18hr and assessed by flow cytometry.

**Figure S6, related to figure 7.**
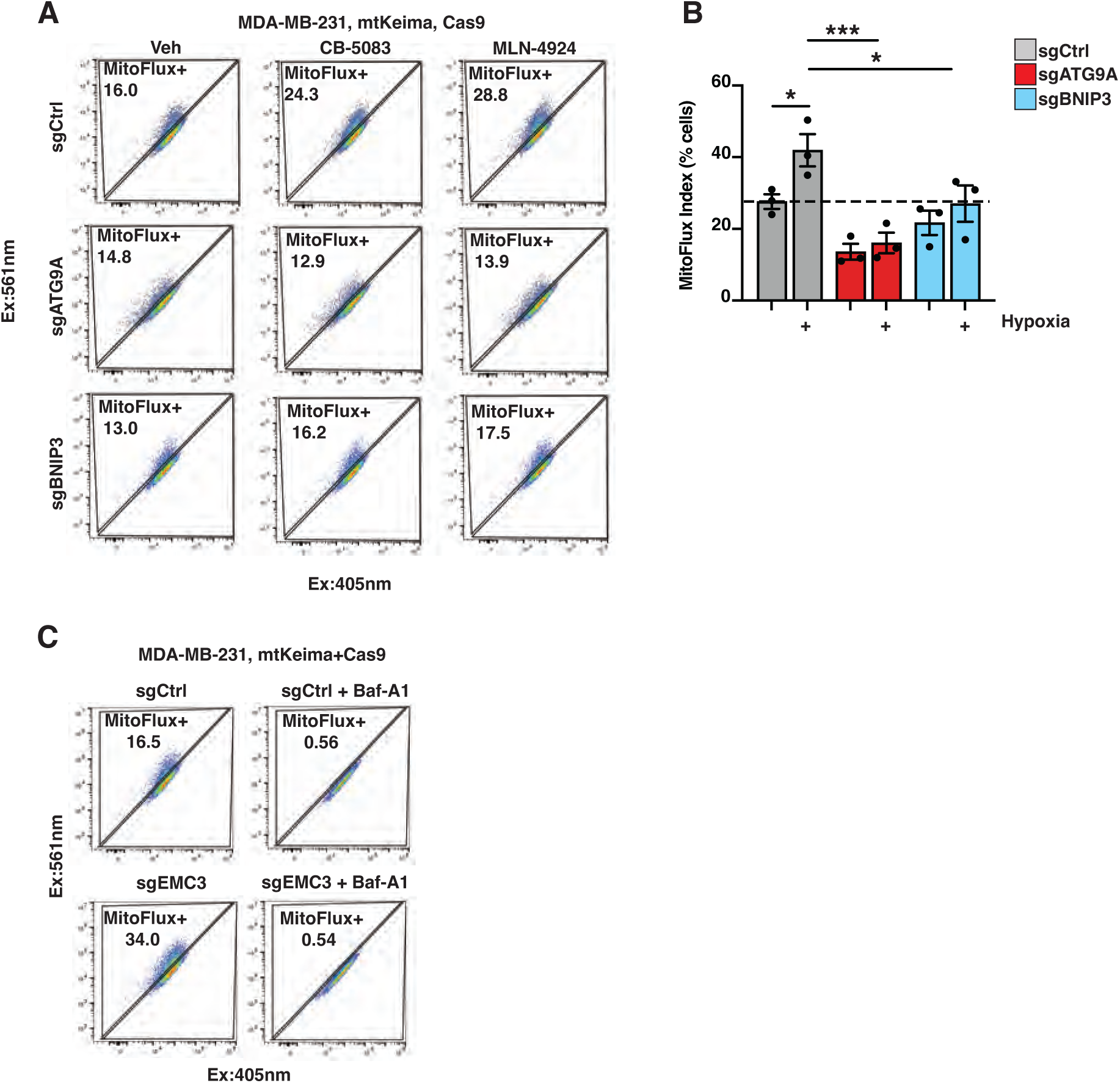
**(A)** MDA-MB-231 mt-Keima cells were transduced with the indicated sgRNAs. On day 8 post-transduction, cells were treated with vehicle (DMSO), MLN-4924 (1*µ*M), and CB-5083 (1*µ*M) for 18hr prior to analysis by flow cytometry. **(B)** MDA-MB-231 cells expressing mt-Keima were transduced the indicated sgRNAs. On day 8 post-transduction, cells were incubated in normoxic and hypoxic conditions for 18hr prior to flow cytometry. Bar graphs represent mean +/- SEM from 3 independent experiments. Statistical analysis was performed using two-way ANOVA with Tukey’s post-test. ***, *p* < 0.001; *, *p* < 0.05. **(C)** MDA-MB-231 cells expressing mt-Keima were transduced with either a non-targeting sgRNA (sgCtrl) or sgEMC3. On day 8 post-transduction, cells were incubated with vehicle (DMSO) or Baf-A1 (100nM) for 18hr prior to flow cytometry.

