## Supplementary material for "The ER membrane protein complex governs lysosomal turnover of a mitochondrial tail-anchored protein, BNIP3, to restrict mitophagy": Table S2

**Table S2: Oligonucleotides**

**Oligonucleotides**

| Sequence | Source | Identifier |
| --- | --- | --- |
| sgRNA targeting sequence: ATG9A Forward: CCGTTTCCAGAACTACATGG | Doench et al. 2016 | Addgene# 73178 |
| sgRNA targeting sequence: BNIP3 Forward: TCTTGTGGTGTCTGCGAGCG | Doench et al. 2016 | Addgene# 73178 |
| sgRNA targeting sequence: ATG7 Forward: CTCTTGTAAATACCATCTGT | Doench et al. 2016 | Addgene# 73178 |
| sgRNA targeting sequence: FIP200 Forward: TTTCTAACAGCTCTATTACG | Doench et al. 2016 | Addgene# 73178 |
| shRNA targeting sequence: Rab7 Forward: ACGTAGGCCTTCAACACAATTCTCGAGAATTGT  GTTGAAGGCCTACGT | Sigma-Aldrich | N/A |
| sgRNA targeting sequence: EMC2 Forward: TGACAGTATAAGTGGTTGTG | Doench et al. 2016 | Addgene# 73178 |
| sgRNA targeting sequence: EMC6 Forward: CCCCGACAGCGCTGACACCG | Doench et al. 2016 | Addgene# 73178 |
| sgRNA targeting sequence: EMC3 Forward: GTAACATAGGCTTAAAACGG | Doench et al. 2016 | Addgene# 73178 |
| sgRNA targeting sequence: GET4 Forward: ACGAGGCGCACCAGATGTAC | Doench et al. 2016 | Addgene# 73178 |
| sgRNA targeting sequence: USO1 Forward: GCTGCTGCAATAACCCACAC | Doench et al. 2016 | Addgene# 73178 |
| sgRNA targeting sequence: SAR1A Forward: ACCAAGATCAAAAGTTGTAA | Doench et al. 2016 | Addgene# 73178 |
| sgRNA targeting sequence: VPS4A Forward: ATGGCCGAATCCAACACCCA | Doench et al. 2016 | Addgene# 73178 |
| sgRNA targeting sequence: MMGT1 Forward: AATATGAACTATACCGTAAC | Doench et al. 2016 | Addgene# 73178 |
| sgRNA targeting sequence: ASNA1 Forward: CTGAAGTGGATCTTCGTCGG | Doench et al. 2016 | Addgene# 73178 |
| sgRNA targeting sequence: PREB Forward: AGGAGCAGGGGCCTCGACAA | Doench et al. 2016 | Addgene# 73178 |
| sgRNA targeting sequence: CAMLG Forward: ACAGCGGACTCGGTCCAGAG | Doench et al. 2016 | Addgene# 73178 |
| sgRNA targeting sequence: WRB Forward: ACGGATAAGCTCAAAACCCA | Doench et al. 2016 | Addgene# 73178 |
| sgRNA targeting sequence: SEC16A Forward: AGAGACACGCTAGAGGACTG | Doench et al. 2016 | Addgene# 73178 |
| sgRNA targeting sequence: HUWE1 Forward: AAGCGCTCAAATCGGACCAA | Doench et al. 2016 | Addgene# 73178 |
| sgRNA targeting sequence: UBA3 Forward: ATGATCCTGAACATATACAA | Doench et al. 2016 | Addgene# 73178 |
| sgRNA targeting sequence: VTA1 Forward: CACAGTAATAAGCCACCACA | Doench et al. 2016 | Addgene# 73178 |
| sgRNA targeting sequence: ATP6V1B2 Forward: TGCTGGGCTACCACACAATG | Doench et al. 2016 | Addgene# 73178 |
| sgRNA targeting sequence: ATP6V1E1 Forward: AGTTCAACATAGAGAAAGGT | Doench et al. 2016 | Addgene# 73178 |
| shRNA targeting sequence: shEMC Forward:  GACATGATGAAAGGGAATGTACTCGAG TACATTCCCTTTCATCATGTC | Sigma Aldrich | N/A |
| shRNA targeting sequence: BNIP3 Forward: GCCTCGGTTTCTATTTATAATCTCGAGATTATAAA  TAGAAACCGAGGC | Sigma-Aldrich | N/A |
